# Which phonetic contrasts recover information lost to aggregate speech scoring in auditory nerve disorders: a computational framework

**DOI:** 10.64898/2026.01.24.701491

**Authors:** Marta Campi, Elie Partouche, Gregory Gerenton, Paul Avan, Clément Gaultier

## Abstract

Auditory nerve disorders, including auditory neuropathy spectrum disorders (ANSD), show distorted patterns of auditory nerve activity despite preserved spectral analysis of sound in the cochlea, producing impaired speech recognition despite normal auditory sensitivity. Distinct pathophysiological mechanisms affecting auditory-nerve activity have been identified in animal models, but current clinical speech tests were not designed to distinguish among them, because they average performance across phoneme categories, collapsing mechanism-specific confusion patterns into a single intelligibility score. Using computational modeling of auditory nerve responses, we simulated four candidate mechanisms and showed that their encoding disruptions cascade into systematic, phoneme-specific recognition failures: brief consonants were severely degraded while sustained vowels were preserved, and each mechanism produced a distinct confusion pattern. Models trained on ANSD-degraded signals developed compensation strategies that generalized to healthy signals, while the reverse did not, and noise training that benefited healthy models harmed ANSD models, mirroring real-world listening difficulty. Because simulation fixes the generating mechanism by construction, we could then measure directly how much mechanism-specific information each behavioral representation preserves. Aggregate word scoring discarded most of the information retained in the full confusion matrix, yet a handful of targeted phonetic contrasts recovered the mechanism. Simulation uniquely enables this analysis because the underlying mechanism is known, a property currently unavailable in patient datasets. Rather than offering direct diagnosis from behavior alone, the framework quantifies what aggregate scoring destroys and identifies the contrasts that best separate mechanisms, a principled route toward phonetically efficient tests that recover mechanistic information current scoring discards.

## 1 Introduction

Hearing loss represents a global health challenge affecting 466 million people worldwide, with projections reaching 900 million by 2050. The associated economic burden exceeds C216 billion annually in Europe alone [1, 2]. Despite hearing aid development, approximately half of individuals who need amplification do not use it, partly due to poor performance in complex acoustic environments [3]. These limitations are particularly evident in Auditory Neuropathy Spectrum Disorders (ANSD), which affect an estimated 1–10% of those with hearing complaints [4, 5] and are associated with minimal benefit from conventional amplification despite preserved peripheral sensitivity. We use ANSD here in a broad sense, to mean any sensorineural hearing loss in which a neural component dominates, rather than as a synonym for cochlear synaptopathy. This encompasses synaptic loss, but also auditory nerve demyelination, as in multiple sclerosis with extensive myelin loss, and nerve injury, as after vestibular schwannoma surgery.

Patients with ANSD exhibit preserved outer hair cell function, as probed by otoacoustic emissions and/or cochlear microphonic responses, yet show abnormal or absent auditory brainstem responses (ABRs) [6, 7], indicating distorted patterns of auditory nerve activity despite normal cochlear filtering. This neural disruption typically manifests as the characteristic syndrome of “hearing without understanding”: patients detect sounds but struggle to comprehend speech, with audiometric thresholds that substantially underestimate their functional communication difficulties [8], though presentation varies widely and some cases may show minimal speech deficits despite measurable neural abnormalities. The finding that ABRs are disrupted suggests impairment of fine timing cues in auditory neuron activity, contrasting with typical cochlear hearing loss that reduces frequency selectivity and elevates detection thresholds. Reported patients show deficits in gap detection, amplitude modulation sensitivity, and speech envelope tracking not explained by their audiometric profiles, attributed to impaired auditory-neuron activity [9, 10]. Other patients with similar complaints of poor speech recognition and indirect evidence of impaired auditory-neuron activity may show no clear audiological pattern, a condition termed ‘hidden hearing loss’ [11, 12]. If different mechanisms disrupt different aspects of auditory-nerve encoding, they should produce different patterns of speech errors, not simply different amounts of intelligibility loss.

Several candidate pathophysiological mechanisms have been identified along the auditory pathway, operating at different sites. This mechanistic diversity is reflected in the marked clinical heterogeneity of ANSD, with patients differing in the severity of their deficits and often fluctuating across days or acoustic conditions [13]. Temporal jitter from auditory nerve demyelination represents one major mechanism: myelin sheaths ensure rapid, synchronized action potential propagation, and demyelination introduces variable conduction delays that scramble timing across frequency channels [14, 15]. Because phase-locking and cross-frequency coincidence detection in the brainstem rely on precisely aligned inputs [16, 17], this jitter disrupts both. Synaptic loss represents a second mechanism, particularly affecting low- and medium-spontaneous rate (SR) fibers that resist saturation and may matter for encoding speech in noise [18, 19]; losing these synapses reduces population coding while sparing the high-SR fibers that mediate quiet threshold detection, which is why pure-tone audiograms can appear normal despite speech-in-noise deficits. Saturation deficits from reduced neural populations represent a third mechanism, compressing dynamic range and preventing level-tolerant spectral representations [20, 21]. Sporadic loss of inner hair cells [22] or dead cochlear zones [23] can produce similar distortions with little threshold elevation, owing to compensation by surviving hair cells or off-frequency listening. We modeled these categories as four perturbation types: temporal jitter in two variants (uniform and scattered) to capture global versus patchy demyelination, and frequency-selective neural loss and saturation as channel elimination and response truncation (Fig. 1A–B). Throughout, these four perturbations are referred to in the text as uniform jitter, scattered jitter, fiber loss, and truncation, and appear in the figures and tables under the labels Uniform Jitter, Scattered Jitter, Loss, and Truncation.

Because these mechanisms disrupt different aspects of auditory encoding, each should shape the pattern of speech errors differently. Speech categories differ systematically in the neural computations they require. Sustained vowels (formant patterns persisting over 50 to 200 ms) can often be recovered through temporal integration, whereas many consonants depend on brief acoustic events (stop bursts and transitions on the order of 5 to 50 ms) that require millisecond-scale neural synchrony. Each mechanism is therefore expected to produce a distinct distribution of phoneme confusions rather than merely a uniform reduction in intelligibility, so that the pattern of errors, not just their overall rate, may carry mechanistic information [24, 25].

Current diagnostic approaches were not designed to identify the mechanism, for two distinct reasons. First, electrophysiological signatures (absent or abnormal ABRs with preserved OAEs) establish that the deficit is neural rather than cochlear, but they operate at the level of synchrony rather than speech: they confirm the presence of ANSD without identifying which pathophysiological mechanism produces it [7]. Even recent work distinguishing synaptopathy from myelinopathy through compound action potentials [26] characterizes the lesion electrophysiologically, leaving open how it translates into the speech errors a patient actually makes; indeed, the very definition of ANSD rests on deteriorated speech perception, with electrophysiological signs only usually accompanying it [8]. Second, speech assessment is built for a different purpose. Identifying the underlying mechanism would have clinical value: the site of lesion already shows prognostic significance for the choice and expected outcome of intervention [8, 27], illustrating that information beyond the binary presence of ANSD can guide management. Standard speech audiometry, however, was designed to quantify intelligibility, not to identify mechanism: it relies on phonetically balanced lists that sample speech sounds representatively to yield an overall measure of how much is understood. In doing so, it reduces responses to a single score, collapsing the multidimensional pattern of errors that might distinguish one mechanism from another [28]. This applies even to phoneme-specific materials. Tests such as the Nonsense Syllable Test [29], the California Consonant Test [30], and similar paradigms can in principle generate phoneme confusion matrices, yet in routine clinical practice they are scored as a single summary value; recovering the full confusion structure would require substantially more repetitions per phoneme than routine testing permits. The problem, however, is not merely practical: two listeners can reach the same aggregate score through entirely different error patterns, the score identical while the confusions that produced it are not. This raises the central question addressed here: which behavioral representation best preserves the information needed to distinguish underlying ANSD mechanisms?

**Figure 1:**
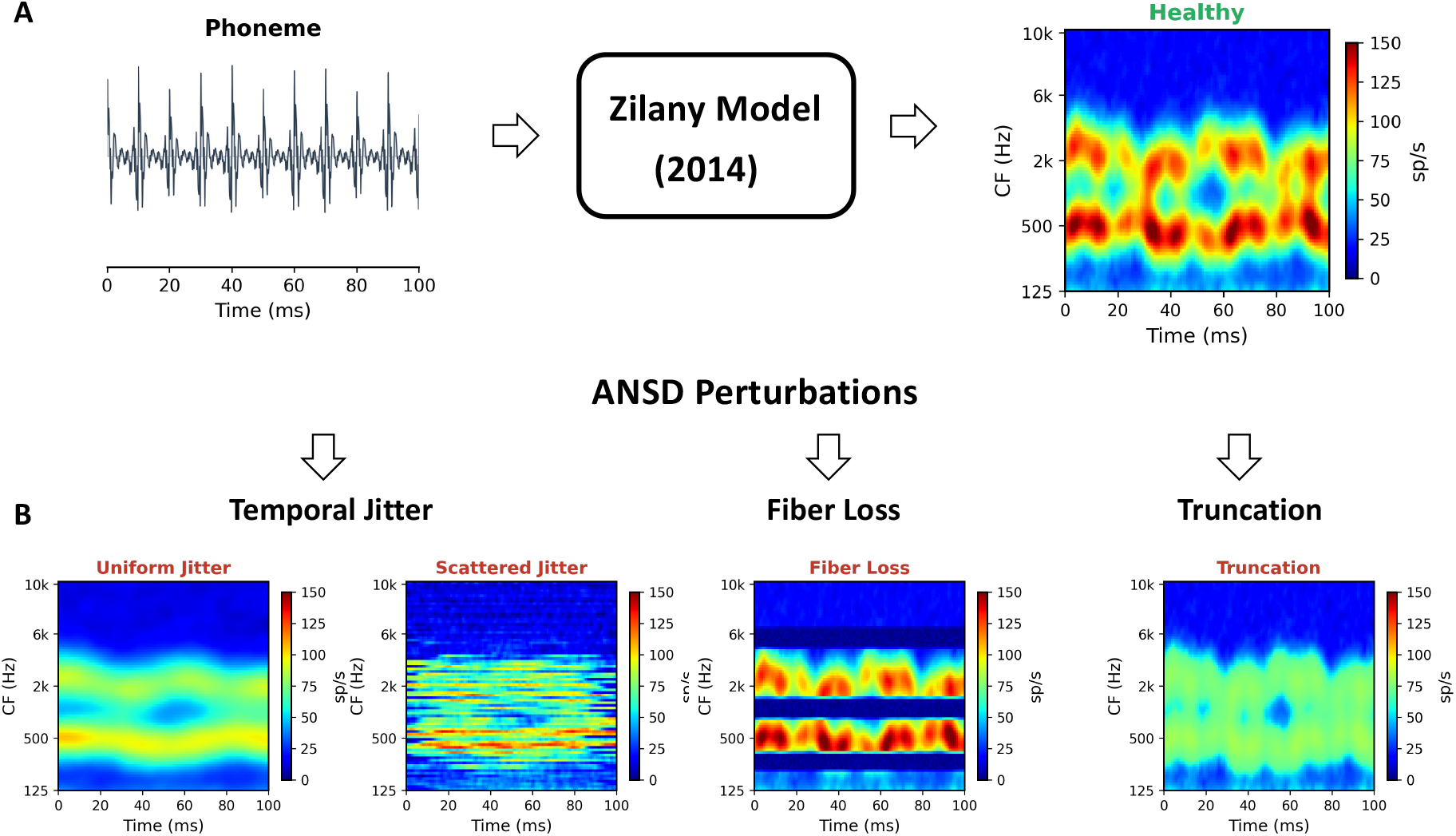
Computational framework for ANSD neurogram perturbations. **(A)** Neurogram generation: speech phonemes are processed through the Zilany auditory nerve model to produce healthy neurograms representing firing rates across characteristic frequency (CF) and time. **(B)** Four perturbation types modeling distinct ANSD mechanisms, each derived from the healthy neurogram. *Temporal jitter* models demyelination in two variants: *uniform jitter* applies identical temporal shifts across all channels, preserving cross-frequency timing; *scattered jitter* applies independent per-channel shifts, destroying cross-frequency synchrony. *Fiber loss* models frequency-selective neural loss (analogous to cochlear dead regions) through channel elimination, creating spectral gaps (dark bands). *Truncation* models neural hypoplasia through amplitude saturation, compressing dynamic range. All panels use identical colorscale (0– 150 sp/s).

Answering this requires known mechanistic ground truth: one must know which mechanism generated each pattern of responses. Although electrophysiological measures can provide indirect evidence for some mechanisms, patient-specific ground truth is generally unavailable, whereas simulation provides it by construction. Only in this setting can competing behavioral representations be evaluated against a known mechanism. To do this, we apply numerical perturbations emulating different ANSD mechanisms to neurograms and feed them to a speech recognition network, exploring all mechanism-phoneme combinations under controlled conditions. This is the approach taken in the present study.

A practical corollary follows for test design. The full confusion matrix is impractical to collect clinically, but it is needed only during the discovery phase: characterizing the complete confusion structure under known mechanisms identifies the small set of phonetic contrasts that an efficient clinical test ultimately needs to measure. This separates a one-time discovery step from the translational test it informs.

## 2 Results

Category-specific confusion patterns then carry mechanism-discriminative information lost to aggregate scoring. We tested this hypothesis in three stages: first quantifying how each proposed ANSD mechanism alters neural encoding of speech (Experiment 1), then determining whether these encoding distortions produce phoneme-specific classification failures (Experiment 2), and finally evaluating whether the resulting patterns persist under realistic noise conditions (Experiment 3). We then asked whether the resulting confusion structure survives the aggregate scoring used in clinical speech audiometry, and whether a small set of targeted contrasts can recover the underlying mechanism (Experiment 4); a further analysis confirms that this holds across closed-set and open-set test formats (SI, Table S17). Testing this hypothesis required constructing three computational modules: a neurogram perturbation generator, a hierarchical speech classifier, and a noise robustness protocol, described below.

### 2.1 Model Construction

Each module described below supports one experiment. Starting from healthy neurograms generated from TIMIT speech stimuli [31] using a biophysically grounded auditory nerve model [32], we introduced four perturbation types simulating proposed ANSD mechanisms (Fig. 1): uniform temporal jitter (identical delays across frequency channels, preserving cross-frequency timing), scattered jitter (independent per-channel delays, disrupting cross-frequency synchrony), selective fiber loss (frequency-channel elimination; labeled “Loss” in figures and tables), and truncation (amplitude saturation).

#### Module 1: Distortion generator

This module quantifies how each perturbation type alters neural encoding of speech using optimal transport theory [33]. The Gromov-Wasserstein distance measures whether phonemes that are acoustically similar remain neurally similar after perturbation. This approach echoes prior work modeling hearing damage effects on neural population responses [34, 35] but targets phoneme-specific disruption patterns rather than aggregate encoding quality. We quantify this structurally, with optimal transport, rather than reading it off a trained classifier’s behavior alone: matching a classifier’s input-output behavior does not by itself establish that the internal mechanism has been captured. A structural measure of how the neural code itself is reshaped therefore complements the behavioral readout the classifier provides in Modules 2 and 3. Encoding disruption does not guarantee perceptual failure: the auditory system might compensate downstream. Experiment 1 tests whether different mechanisms produce distinct category-specific encoding disruptions.

#### Module 2: Classifier

Building on [36], this module comprises a two-stage hierarchical speech recognition system: Stage 1 extracts acoustic-phonetic features from brief neurogram windows (100 ms), Stage 2 integrates predictions across longer temporal context (610 ms), roughly paralleling early auditory feature extraction and cortical integration. The two-stage design can reveal whether ANSD failures arise at the encoding level, the integration level, or from their interaction. The module also includes a misclassification analysis approach inspired by reverse correlation methods [37, 38, 39], averaging neurograms conditioned on classification outcome to reveal how perturbations physically transform the signal for specific phonemes. Experiment 2 tests whether encoding distortions translate to systematic classification failures.

#### Module 3: Noise robustness

This module evaluates robustness by testing individual perturbation types in isolation to establish mechanism-specific severity rankings, and by introducing environmental noise at varying signal-to-noise ratios to compare silence-trained and noise-trained models. Experiment 3 tests whether category-specific patterns persist under realistic listening conditions.

### 2.2 Experiment 1: Each Mechanism Distorts Specific Phoneme Categories

Optimal transport analysis quantified how ANSD perturbations alter the correspondence between acoustic formant trajectories and neural representations. The Gromov-Wasserstein (GW) distance measures whether this correspondence is preserved: higher distances indicate the neural code no longer maintains acoustic similarity structure; sounds that should be encoded similarly become neurally dissimilar. Concretely, GW distance quantifies whether phonemes that are acoustically similar (close formant trajectories) remain neurally similar after perturbation: a distance of zero indicates perfect preservation, while higher values indicate that perturbations have scrambled the relationship between acoustic and neural similarity.

Overall GW distances differed significantly across the four perturbation types (Kruskal-Wallis *H* = 10.02, *p* = 0.018; Fig. 2A). However, this between-mechanism difference is small compared to variation within mechanisms: different phoneme categories showed nearly 4-fold variation in GW distance under the same perturbation type. That is, phoneme-specific patterns carry greater diagnostic information than aggregate mechanism severity.

We analyzed GW distances both across phoneme categories and across individual formant trajectories (F1, F2, F3) to characterize category-specific and frequency-specific disruption patterns.

#### Category-specific patterns provide diagnostic signatures

Different ANSD mechanisms produced divergent phoneme profiles enabling mechanism identification (Table 1; Fig. 2C).

##### High-disruption categories

Fricatives, glottal stops, and flaps consistently showed highest GW distances. These categories are particularly affected by temporal disruption due to their brief durations, aperiodic components, and rapid spectral transitions. Flaps showed peak disruption with scattered jitter (0.070), consistent with their requirement for cross-frequency synchrony to detect the brief 20–30 ms ballistic gesture.

##### Low-disruption categories

Vowels, nasals, and liquids maintained low distances (0.018–0.037) across all perturbations. Their sustained, quasi-periodic structure can be extracted despite timing imprecision through temporal integration.

##### Perturbation-specific signatures

Uniform jitter disproportionately affected fricatives (0.052) and glottal stops (0.070), 58% higher fricative disruption than scattered jitter (0.033). This reflects uniform jitter’s preservation of cross-frequency relationships: fricative identification depends on within-channel spectral shape, which uniform jitter degrades by smearing temporal structure. Scattered jitter produced the most severe flap disruption (0.070), the highest single category-perturbation combination, because flap detection requires cross-frequency coincidence that scattered jitter destroys. Fiber loss perturbation showed highest impact on affricates (0.052), which require intact encoding across multiple frequency regions. Truncation showed most uniform effects across categories (SD = 0.015), consistent with dynamic range compression affecting all phonemes proportionally.

**Table 1:** Gromov-Wasserstein distances (dimensionless, relative scale) quantifying encoding disruption between acoustic formant trajectories and neural representations, by phoneme category and perturbation type. The Healthy column gives the baseline acoustic-neural distance with no perturbation applied; perturbation values are read as increases above this intrinsic baseline. Higher values indicate greater distortion of acoustic-neural correspondence. Values range from 0.018 (Vowel, Truncation) to 0.083 (Flap, Uniform).

| Category | Healthy | Uniform | Scattered | Loss | Trunc |
| --- | --- | --- | --- | --- | --- |
| Vowel | 0.005 | **0.028** | 0.023 | 0.021 | 0.018 |
| Nasal | 0.007 | **0.034** | 0.021 | 0.033 | 0.025 |
| Liquid | 0.006 | **0.037** | 0.033 | 0.033 | 0.032 |
| Fricative | 0.008 | **0.052** | 0.033 | 0.049 | 0.037 |
| Stop | 0.007 | 0.042 | 0.029 | 0.048 | **0.065** |
| Affricate | 0.006 | 0.051 | 0.037 | **0.052** | 0.047 |
| Flap | 0.009 | **0.083** | 0.070 | 0.071 | 0.068 |
| Glottal | 0.008 | 0.070 | 0.054 | 0.064 | **0.076** |

#### Formant-specific patterns

Higher formants showed greater disruption (Fig. 2B): F3 (∼ 2500 Hz) exhibited GW distances of 0.046–0.058 across perturbations, while F1 (∼ 500 Hz) showed lowest distances (0.035–0.039). This gradient reflects differential temporal demands: F1 tracks slow jaw movements (5–10 Hz modulations) resilient to timing imprecision; F3 encodes rapid articulatory gestures (15–25 Hz) where millisecond-scale jitter becomes comparable to modulation periods. Uniform jitter showed the steepest gradient (66% F1-to-F3 increase), while scattered jitter produced more uniform disruption across formants (24% gradient) by destroying both within-channel timing and cross-channel synchrony.

However, the 4-fold category variation in encoding distortion predicts that classification should fail differently across phoneme categories. Whether this prediction holds requires testing speech recognition directly.

### 2.3 Experiment 2: Encoding Distortions Produce Systematic Classification Failures

Experiment 1 established encoding distortions; Experiment 2 tests whether these translate to classification failures. We proceed in two logical steps: matched testing first, where a model trained on one population is tested on that same population, then cross-population transfer, which asks whether the features a model has learned still suffice when it receives signals from the population it was not trained on. If ANSD destroys temporal structure, integration that benefits healthy listeners may harm ANSD listeners by accumulating corrupted features rather than compensating for them. Cross-population transfer tests whether learned strategies remain adequate when the classifier receives neurograms from the population it was not trained on: if healthy-trained models fail on ANSD signals, they depend on features that ANSD destroys; if ANSD-trained models succeed on healthy signals, they have learned to exploit features that survive temporal degradation.

**Figure 2:**
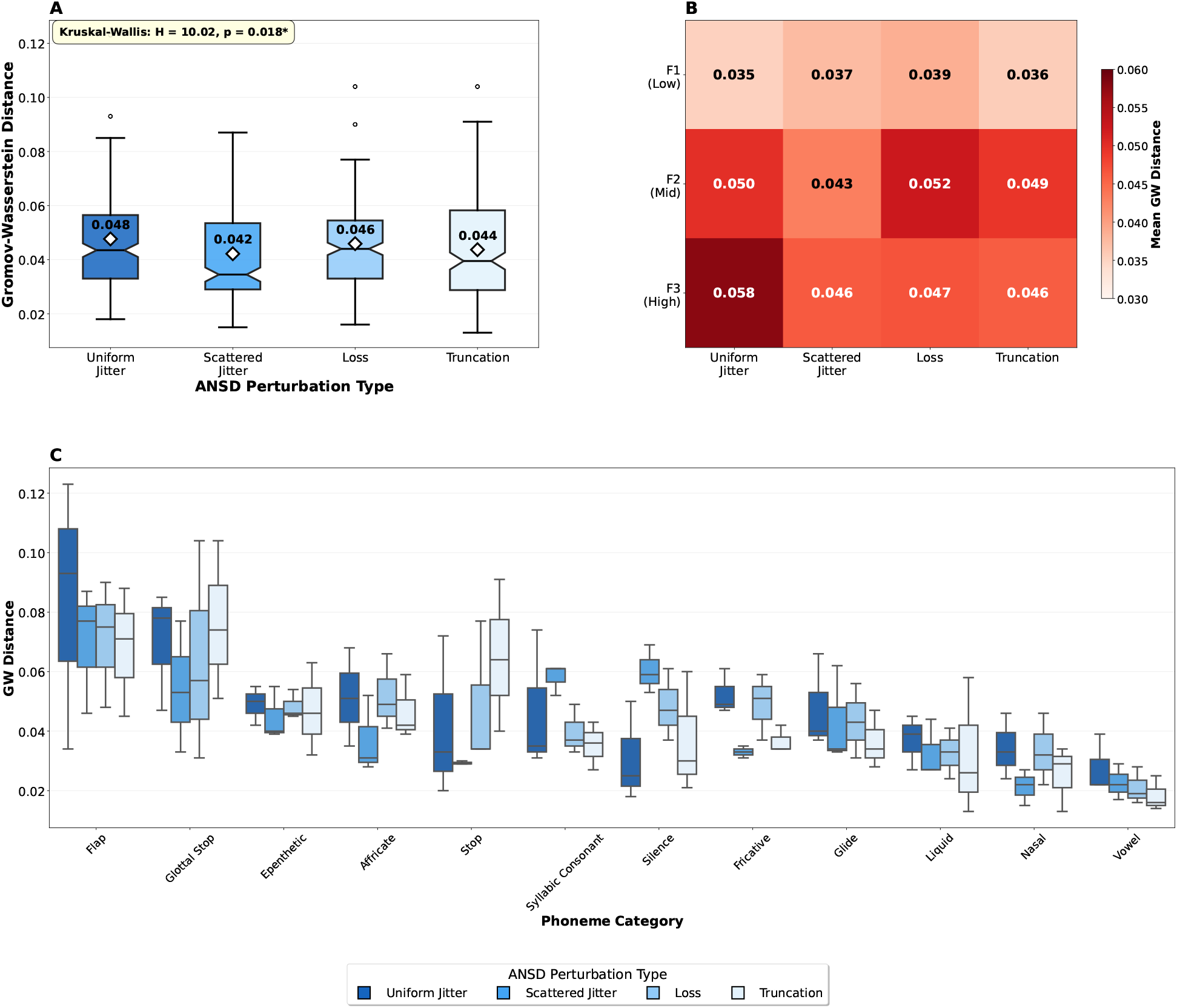
Experiment 1: Optimal transport reveals phoneme-specific encoding disruptions. **(A)** Overall perturbation effects show significant differences across ANSD mechanisms (Kruskal-Wallis *H* = 10.02, *p* = 0.018). White diamonds and the annotated values indicate means; boxes show the interquartile range around the median. Perturbation order: Uniform Jitter, Scattered Jitter, Loss, Truncation. **(B)** The same means decomposed by formant (mean GW distance, as in Panel A): distances increase with formant frequency under jitter, most steeply for uniform jitter (66% from F1 to F3), whereas loss and truncation peak at F2. **(C)** Category-specific distribution showing diagnostic signatures across perturbation types: flaps are most sensitive to uniform jitter and glottal stops to truncation, whereas vowels and nasals remain preserved across all perturbations; per-category means are reported in Table 1. Minor categories (epenthetic, syllabic consonant, silence, glide) are shown for completeness but not discussed in the text.

We trained four models to disentangle population and environmental effects. Two models were trained on healthy (unperturbed) neurograms: Healthy-Silence and Healthy-Noise. Two models were trained on ANSD-perturbed neurograms: ANSD-Silence and ANSD-Noise. ANSD training used mixed perturbations, each utterance received one randomly sampled perturbation type (fiber loss, truncation, uniform jitter, or scattered jitter), to simulate clinical heterogeneity and ensure models learned general rather than mechanism-specific strategies. Noise-trained models incorporated additive environmental noise at SNRs uniformly sampled from 0–20 dB.

This design enables three key analyses: matched testing, cross-population transfer, and misclassification analysis. Robustness testing (Experiment 3) examines noise interactions and mechanism-specific effects. All results use TIMIT’s held-out test set (168 speakers, no overlap with training). All category-specific accuracy differences were significant (McNemar’s test, *p <* 0.001; bootstrap 95% CIs in SI Tables S6–S8).

#### Matched testing reveals hierarchical failure and category-specific patterns

Both populations showed hierarchical change, the difference between Stage 1 and Stage 2 accuracy, when tested on held-out speakers, but the magnitude differed significantly (Fig. 3A). Healthy-Silence models showed modest Stage 2 reduction (Stage 1: 67.0% to Stage 2: 61.0%). ANSD-Silence models showed severe degradation (48.0% to 25.5%), a 22.5-point loss, nearly four times the healthy reduction.

**Figure 3:**
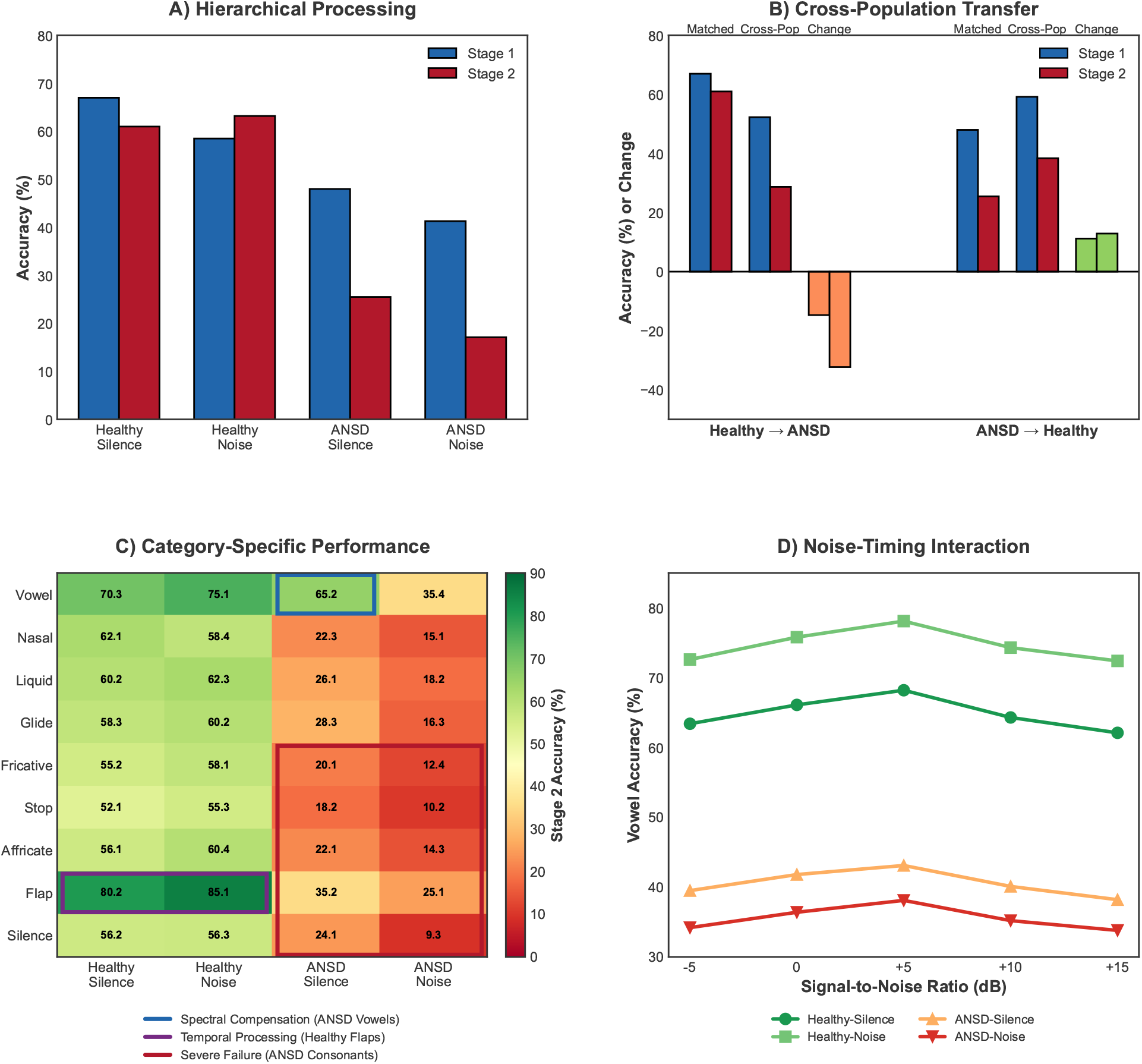
Speech recognition reveals hierarchical failure, asymmetric transfer, and noise interactions (Experiments 2 and 3). Each population is shown in both silence and noise. **(A)** Hierarchical processing (Stage 1 versus Stage 2) on held-out speakers, for Healthy and ANSD models in silence and in noise: Healthy models show a modest Stage 2 change, whereas ANSD models degrade severely, and noise training widens this gap. **(B)** Cross-population transfer in silence (matched, cross-population, and their change): Healthy models fail on ANSD data, whereas ANSD models improve on healthy data. The corresponding noise results are reported in the text (Experiment 3). **(C)** Category-specific Stage 2 accuracy across the phoneme inventory, for Healthy and ANSD models in silence and noise: ANSD preserves vowel discrimination (65.2%, blue box) but shows severe consonant failure (red box), while Healthy flaps retain intact temporal processing (purple box). **(D)** Noise-timing interaction: vowel accuracy across signal-to-noise ratios for silence-trained and noise-trained Healthy and ANSD models; noise training benefits Healthy but harms ANSD.

ANSD-Silence models achieved strong vowel accuracy (65.2%), approaching healthy performance (70.3%) and far exceeding other categories (Fig. 3C; SI Table S6 for all categories). Temporal consonants showed severe impairment that hierarchical processing could not rescue. Stops achieved only 18.2% (vs. 52.1% healthy), fricatives 20.1% (vs. 55.2%), and flaps 35.2% (vs. 80.2%). The 45-point flap deficit represents the largest category-specific gap, consistent with flaps’ brief duration (20–30 ms) requiring precise cross-frequency synchrony.

Confusion matrices revealed systematic rather than random errors (Table 2). Flaps were misclassified as vowels 26.3% in ANSD versus 1.8% healthy (14.6-fold increase). Stops showed 22.1% vowel confusion (vs. 3.2%), fricatives 24.3% (vs. 2.1%; complete confusion matrices in SI Fig. S1).

#### Cross-population transfer reveals fundamental asymmetry

Cross-population testing revealed striking asymmetries (Fig. 3B).

Healthy models failed severely when tested on ANSD data: Stage 2 accuracy dropped from 61.0% (matched) to 28.7% (transfer) in silence, a 32-point loss. Remarkably, ANSD models *improved* when tested on healthy data, rising from 25.5% (matched) to 38.4% (transfer), a 13-point gain.

This asymmetry is unidirectional: strategies learned on degraded signals generalize to intact signals, but not the reverse.

#### Misclassification analysis reveals a stop-to-vowel transformation mechanism

To identify spectrotemporal features driving the consonant-to-vowel confusions documented above, we computed classification-conditioned templates: average neurograms for tokens grouped by predicted category and correctness. Because the classifier learns decision boundaries in neural space, averaging neurograms conditioned on classification outcome recovers the prototypical pattern underlying each decision. Comparing correctly and incorrectly classified templates therefore isolates the spectrotemporal changes that drive representations across category boundaries.

We focused on stop /t/ (canonical high-frequency burst, prominent in confusions) and vowels /aa/, /ih/, /iy/ spanning distinct formant configurations (additional consonants in SI Fig. S5; perturbation-specific templates in SI Fig. S3).

When healthy models correctly classified /t/, templates showed intense, brief broadband bursts (0–25 ms, 2–10 kHz). When ANSD models misclassified /t/ as vowel (22.1% of tokens), templates showed reduced early burst energy (0– 50 ms) accompanied by sustained activity extending 50–100 ms into formant-frequency regions (Fig. 4A).

**Figure 4:**
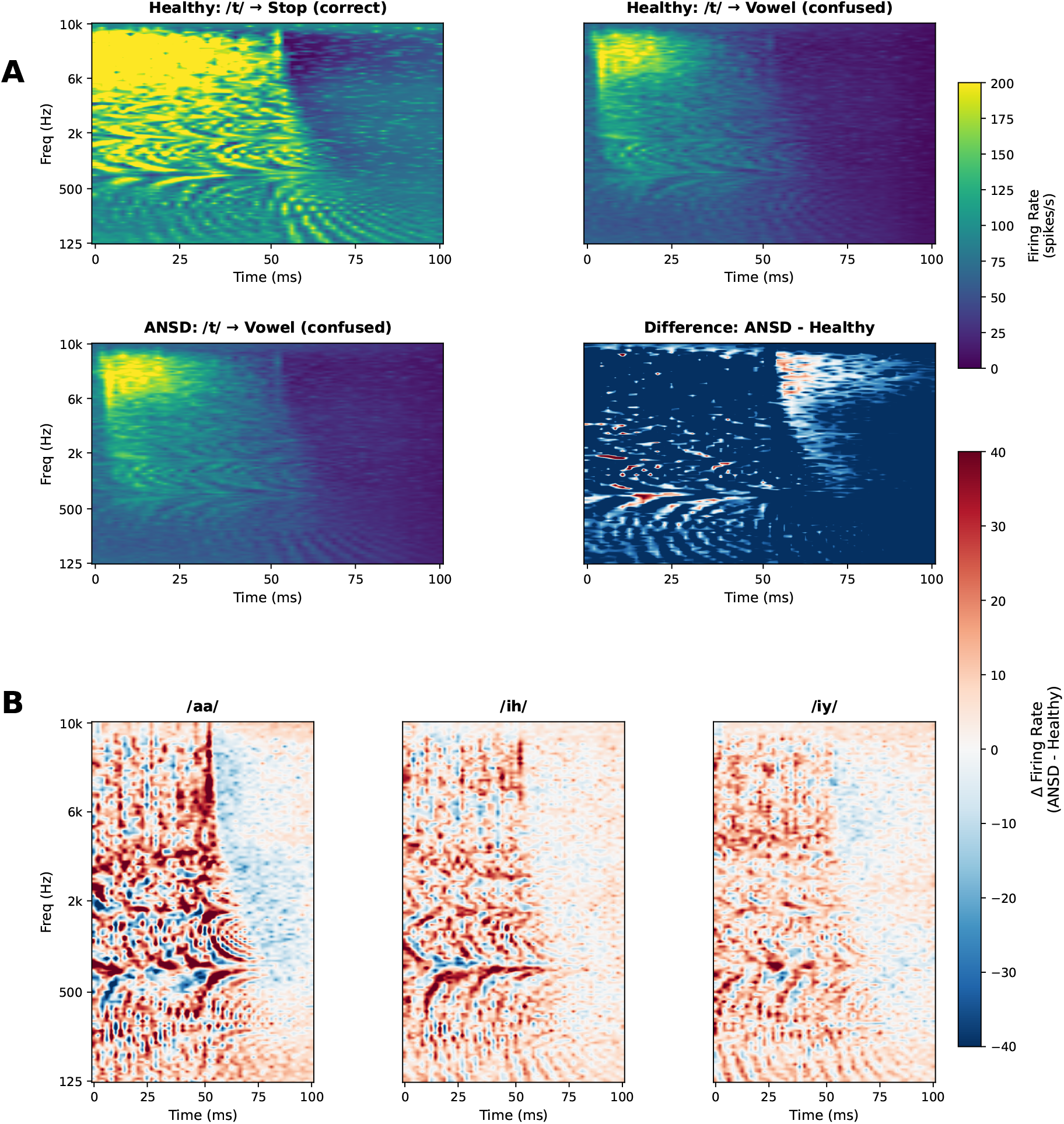
Misclassification analysis reveals a stop-to-vowel shift in neural representation. **(A)** Stop /t/ templates separated by population and classification outcome. Healthy correct classifications (top left) show the canonical brief high-frequency burst (0–25 ms, 6–10 kHz). When classified as vowels, healthy tokens (top right) exhibit attenuated burst structure. In ANSD, tokens confused as vowels (bottom left) show marked reduction of early burst energy accompanied by increased sustained low-frequency activity (50–100 ms, 500–3000 Hz). ANSD confused tokens closely resemble healthy confused tokens, indicating that temporal perturbations shift consonant representations toward neural regions associated with vowel decisions, rather than generating qualitatively new error patterns. The difference map (bottom right; ANSD confused minus Healthy correct) quantifies this redistribution: blue regions (0–50 ms) indicate diminished transient burst energy, whereas red regions (50–100 ms, formant frequencies) indicate enhanced sustained activity. **(B)** Vowel difference maps (/aa/, /ih/, /iy/) show ANSD-induced distortions (red: low-frequency excess below 2 kHz; blue: mid-frequency reduction) but preservation of overall spectral organization, consistent with retained vowel classification.

This consonant-to-vowel bias was not limited to /t/ but generalized across manner classes (Table 2).

**Table 2:**
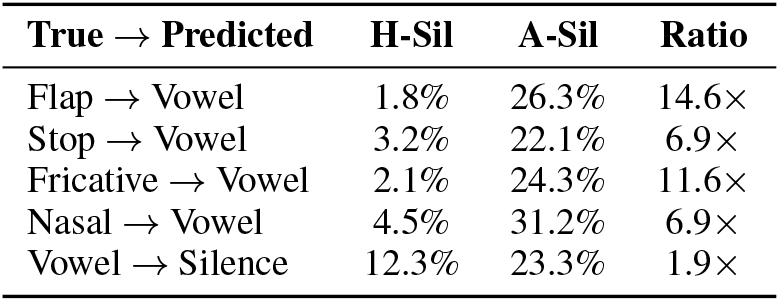
Systematic confusion patterns showing consonant-to-vowel transformation in ANSD. H-Sil: Healthy-Silence; A-Sil: ANSD-Silence. Ratio indicates fold-increase in ANSD relative to healthy. Flaps show greatest transformation (14.6*×*), consistent with their brief duration and cross-frequency synchrony requirements.

| True → Predicted | H-Sil | A-Sil | Ratio |
| --- | --- | --- | --- |
| Flap → Vowel | 1.8% | 26.3% | 14.6× |
| Stop → Vowel | 3.2% | 22.1% | 6.9× |
| Fricative → Vowel | 2.1% | 24.3% | 11.6× |
| Nasal → Vowel | 4.5% | 31.2% | 6.9× |
| Vowel → Silence | 12.3% | 23.3% | 1.9× |

The difference map (ANSD confused minus Healthy correct) revealed a redistribution of energy from early high-frequency transients (blue, 0–50 ms) to later low-frequency sustained structure (red, 50–100 ms, 500–3000 Hz), consistent with a shift from stop-like to vowel-like representation.

Importantly, healthy confused and ANSD confused templates were highly similar (Fig. 4). ANSD does not generate a novel error pattern; rather, temporal perturbations systematically shift consonant representations into neural regions already associated with vowel decisions.

Vowel templates (/aa/, /ih/, /iy/) showed ANSD-induced distortions including high-frequency reduction and altered formant structure, yet preserved sufficient spectral organization for reliable classification (Fig. 4B).

### 2.4 Experiment 3: Mechanism-Specific Patterns Persist in Noise

Experiment 2 established category-specific failures using silent training conditions. Experiment 3 tested robustness: how noise training interacts with temporal disruption, whether deficits persist across SNR levels, and which individual perturbation mechanisms pose the greatest challenges. If ANSD simply adds internal noise, noise training should help both populations equally by teaching robustness to stochastic variability. Divergent responses would indicate that ANSD involves structural signal degradation, not just increased variability.

#### Noise-timing interaction

Noise training and temporal disruption interact: the same training regime that benefits healthy populations harms ANSD populations (Fig. 3A,D). Healthy models showed hierarchical benefit under noise (Stage 1: 58.5% to Stage 2: 63.2%), while ANSD models showed further degradation (41.3% to 17.1%). Noise training improved healthy vowel accuracy (from 70.3% to 75.1%) but degraded ANSD vowel accuracy (from 65.2% to 35.4%), a 30-point drop.

This opposite effect demonstrates that ANSD does not simply add internal noise. Noise training reinforces temporal processing strategies that ANSD signals cannot support, which may explain why the performance gap widens under noise (35.5 points in silence to 46.1 points in noise).

The transfer asymmetry confirmed this pattern: healthy-trained models’ failure on ANSD data grew from − 32.3 points in silence to − 43.8 under noise, while ANSD-trained models’ success on healthy data remained stable (+12.9 to +13.5 points; SI Table S12).

#### Persistent deficits across SNR levels

Testing across signal-to-noise ratios (+15 to − 5 dB) confirmed the Healthy-ANSD performance gap persists regardless of environmental noise level (Fig. 3D). The gap remains approximately constant across all SNRs. Noise training provided consistent benefit to healthy vowel accuracy (+10 points across SNRs) but consistently harmed ANSD vowel accuracy (−5 points; SI Table S13).

#### Mechanism-specific severity hierarchy

The preceding analyses used mixed-perturbation training and testing. To isolate effects of individual ANSD mechanisms, the same four models were tested on single perturbation types applied in isolation (Table 3; see SI Table S11 for complete category-by-category accuracy breakdowns). Testing healthy models on perturbations provides a baseline; comparing to ANSD models reveals which mechanisms benefit most from adapted training.

This established a severity ranking (Table 3): fiber loss *<* truncation *<* uniform jitter *<* scattered jitter. Fiber loss perturbation uniquely showed hierarchical benefit across all four populations, the only perturbation where Stage 2 exceeded Stage 1. Temporal disruptions showed negative hierarchical Δ, with scattered jitter most severe (SI Table S11 for category breakdowns; statistical validation in SI Table S10).

### 2.5 Experiment 4: Aggregate Scoring Discards Mechanism Information That Contrasts Recover

Experiments 1 to 3 established that different ANSD mechanisms produce distinct patterns of phoneme confusions. Experiment 4 asks a different question: if these same responses are summarized the way clinical speech tests summarize them, is the underlying mechanism still distinguishable? The Stage 2 confusion matrices serve two complementary purposes. They first provide the mechanism-specific confusion patterns directly, read from the matrices as they are, without sampling. They are then used as probabilistic models of virtual listeners: each confusion matrix specifies, for every presented phoneme category, the probability of each perceived phoneme category, and sampling from these probabilities generated a response to every phoneme in the held-out TIMIT sentences, yielding complete synthetic responses whose underlying mechanism is known by construction. We then summarized these same responses as progressively simpler behavioral representations, from the full pattern of confusions down to the single word-recognition score a clinic records, and measured how much information about the underlying mechanism survives each step (Methods).

#### The mechanisms differ in confusion structure even when their scores converge

A single overall score, the mean of each mechanism’s confusion-matrix diagonal (the number a clinic records), separates the four mechanisms only by degree: fiber loss stands apart (51.7% correct), while the three temporal mechanisms cluster together (truncation 25.7%, jitter 21.1%, scattered 22.5%). Where their errors go, however, differs sharply. Fiber loss and jitter send most consonant errors to vowels (72% and 73%), whereas truncation sends far fewer to vowels (32%) and instead confuses consonants with one another (44% obstruent-to-obstruent, among fricatives, stops, and affricates); scattered jitter is intermediate. The consequence is clearest for truncation and jitter: the two differ by only 4.6 pp in overall accuracy, yet their confusions among the obstruent categories differ by 18 to 22 pp, and in the vowel column by 10 to 13 pp (Fig. 5A). An aggregate score treats them as nearly identical; their error structure places them far apart.

**Table 3:**
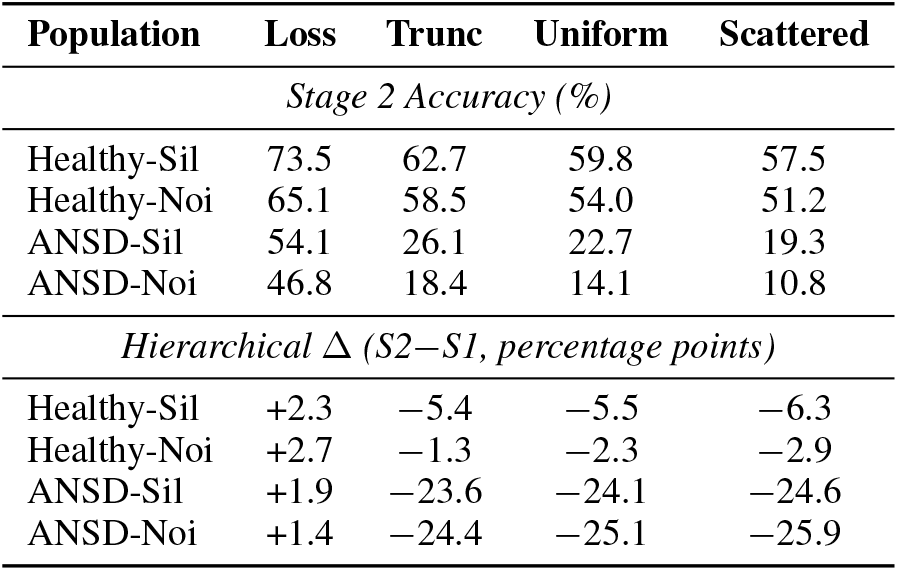
Mechanism-specific severity hierarchy. Models trained with mixed perturbations (ANSD) or healthy neurograms (Healthy) were tested on individual perturbation types applied in isolation. Stage 2 accuracy and hierarchical change (S2 − S1) shown. Loss uniquely preserves hierarchical benefit across all populations; temporal perturbations produce failure, with scattered jitter most severe. Complete category-by-category accuracy breakdowns in SI Table S11.

| Population | Loss | Trunc | Uniform | Scattered |
| --- | --- | --- | --- | --- |
| *Stage 2 Accuracy (%)* |  |  |  |  |
| Healthy-Sil | 73.5 | 62.7 | 59.8 | 57.5 |
| Healthy-Noi | 65.1 | 58.5 | 54.0 | 51.2 |
| ANSD-Sil | 54.1 | 26.1 | 22.7 | 19.3 |
| ANSD-Noi | 46.8 | 18.4 | 14.1 | 10.8 |
| *Hierarchical $\Delta$ (S2–S1, percentage points)* |  |  |  |  |
| Healthy-Sil | +2.3 | –5.4 | –5.5 | –6.3 |
| Healthy-Noi | +2.7 | –1.3 | –2.3 | –2.9 |
| ANSD-Sil | +1.9 | –23.6 | –24.1 | –24.6 |
| ANSD-Noi | +1.4 | –24.4 | –25.1 | –25.9 |

**Figure 5:**
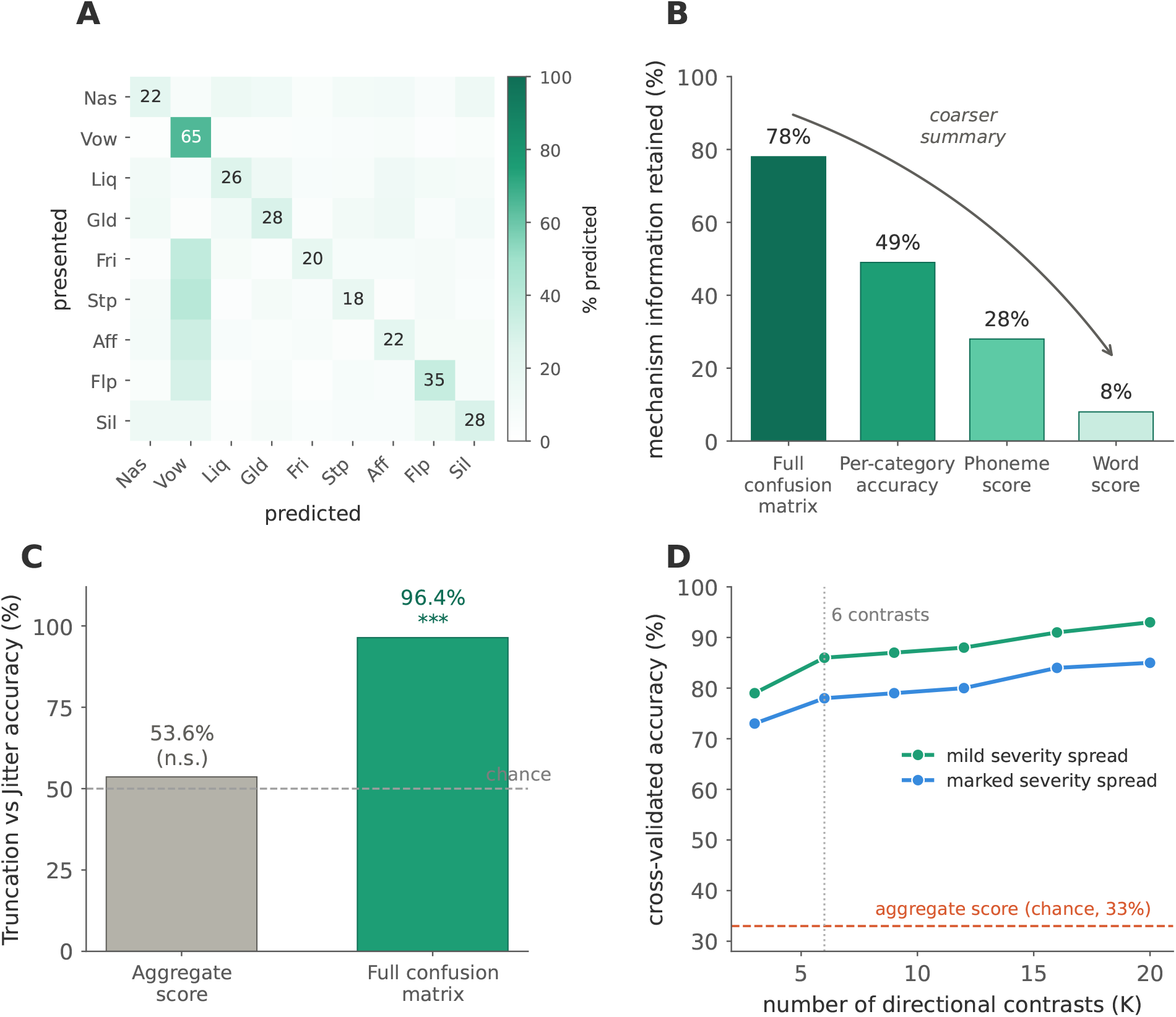
Aggregate scoring destroys mechanism-discriminative information that the confusion-matrix structure preserves, and a few contrasts recover. **(A)** Reference ANSD confusion-matrix structure (Stage 2): rows are the true phoneme category, columns the predicted category, and the diagonal the proportion correct. The mechanism-specific differences in error routing between truncation and jitter are quantified in the text. **(B)** Mutual information about the mechanism retained at each level of aggregation, from the full confusion matrix down to the word score, where only 8% survives. **(C)** A nearest-centroid classifier separates truncation from jitter at chance from the aggregate score (53.6%) but almost perfectly from the full confusion matrix (96.4%). **(D)** Cross-validated classification of the three temporal mechanisms as a function of the number of directional contrasts, under a realistic listener model (inter-listener variability, variable severity, variable item counts): accuracy rises and saturates near 79 to 86% from six contrasts, against the 33% chance level of the aggregate score (dashed). The five scoring models, the cross-transfer and open-set (VCV) analyses, and the Gromov-Wasserstein structural distance are reported in the SI (Tables S14, S15, S17, S18).

#### No scoring of the responses recovers the difference

None of the five scoring rules recovered the distinction between the temporal mechanisms. Across increasingly permissive rules, from requiring every phoneme correct to allowing one or two errors per word, relaxing the scoring raised recognition scores for all mechanisms but never separated the temporal ones, whose scores stayed within 5.3 pp of one another (SI, Table S14). Only fiber loss remained consistently distinguishable.

This result proved robust across several further analyses (SI, Tables S15 and S17). It held whether or not the simulated listener’s compensation strategy matched its pathology (for example, a normally developed auditory system with sudden ANSD onset; SI, Table S15), and whether the test was closed-set or open-set. None of these restored separation among the temporal mechanisms: the convergence does not depend on the particular listener model or test format.

#### Aggregation discards most of the mechanism information

We quantified this loss by representing the same responses at progressively coarser levels. As the summary coarsens from the full confusion matrix down to the single word score a clinic records, the mutual information between mechanism and responses, that is, how much the responses reveal about which mechanism produced them, collapses steadily: the full matrix retains 78% of the available information, per-category accuracy 49%, the phoneme score 28%, and the clinical word score only 8% (Fig. 5B; Methods). Word-level scoring therefore discards 92% of the mechanism information the confusion matrix preserves. A simple classifier that identifies the mechanism from the responses confirms this: it does no better than chance when given only the aggregate score (53.6%), but is almost always correct when given the full confusion matrix (96.4%; Fig. 5C). This is not an artifact of how the information was estimated, since a bound that does not rely on that estimation gives the same picture (Methods). The mechanisms differ in how they make errors, not in how many errors they make. Aggregation preserves the latter while discarding the former.

#### A few targeted contrasts recover the mechanism

The information that aggregate scoring discards is not spread across the confusion matrix; it is concentrated in a small number of directional contrasts. Ranking the matrix cells by how differently they behave across the three temporal mechanisms identifies only a handful of such contrasts. Using these alone, a nearest-centroid classifier separated the three temporal mechanisms well above chance: six directional contrasts were enough to reach 79 to 86% accuracy under five-fold cross-validation, whereas the aggregate score stayed near chance (≈ 33%; Fig. 5D). The contrasts are interpretable, each an error one mechanism makes but another does not: truncation confuses consonants with one another (for example Aff → Flp, Stp → Fri), whereas jitter sends consonants toward vowels and sonorants (for example Stp → Vow, Flp → Nas).

This was not an artificially easy classification problem. Simulated listeners were not assumed to share an identical confusion matrix: each received an individual matrix combining three realistic sources of variability, inter-listener variation, a variable severity level, and a variable number of test items per category (Methods). The contrasts and the class centroids were selected on the training folds only, so contrast selection never saw the test data. Two controls confirm the effect reflects real structure: permuting the mechanism labels collapsed accuracy to chance, and listeners simulated with mixed mechanisms were assigned to one of their two constituent mechanisms rather than to the unrelated third. Aggregate scores discard these contrasts; targeted contrasts recover them.

## 3 Discussion

The central contribution of this work is a computational framework that quantifies how much mechanistic information is lost when speech responses are reduced to aggregate scores. Aggregation always discards something; what has been missing is a way to measure how much. Simulation supplies this by fixing the generating mechanism, a property unavailable in patient datasets, so that competing behavioral representations can be compared directly against the mechanism that produced them. Aggregate word scoring discarded 92% of the mechanism information the confusion matrix preserved, yet a handful of targeted contrasts recovered it (Experiment 4). This changes the question from “can mechanisms be distinguished?” to “which behavioral representation preserves the information needed to distinguish them?”

### Timing survival out of stage 1 determines stage 2 outcomes

Underlying the mechanistic results is a single constraint: once information is discarded by an upstream representation, later stages cannot recreate it. Models trained on ANSD signals learn timing-independent features that generalize to intact signals; models trained on healthy signals learn timing-dependent features that fail once timing is degraded. Stage 2 integrates whatever Stage 1 provides, so misaligned features are amplified rather than corrected. The mechanism-specific breakdown confirms this: only fiber loss preserves hierarchical benefit (+1.9 pp), because timing survives within the channels that remain, while the three temporal perturbations all produce severe hierarchical failure (−23.6 to −24.6 pp; Table 3).

### Phoneme-specific signatures

The same constraint explains why mechanisms disrupt different phonemes. Optimal transport showed that variation across phoneme categories within a single mechanism was nearly four-fold, far exceeding the variation between mechanisms (Experiment 1), so phoneme-level signatures carry more diagnostic information than any aggregate measure. The pattern is mechanistically sensible: uniform jitter degrades fricatives by smearing the within-channel timing their spectral shape depends on, scattered jitter destroys the cross-frequency coincidence that brief flaps require, fiber loss most affects affricates yet preserves hierarchical benefit because timing survives in the remaining channels, and truncation acts uniformly, consistent with dynamic-range compression. These distortions propagated from encoding into classification and persisted in noise (Experiments 2 and 3).

This constraint also predicts opposite effects of noise training in the two populations, and this is what we observe (Fig. 3A): healthy models gain hierarchical benefit under noise, whereas ANSD models worsen further. This dependence on how the representation adapts, rather than on any fixed feature, converges with cortical evidence that encoding flexibility, not the magnitude of any one feature, predicts comprehension of degraded speech [40]. Experiment 4 shows the same principle at the level of behavioral summary: once the confusion structure is compressed into a single score, the discarded mechanistic information cannot be reconstructed.

### Convergence with prior work

Our findings align with recent mechanistic studies: synaptopathy degrades temporal envelope coding [41], spontaneous-rate-specific fiber loss degrades word recognition in Zilany-based simulations [42], and limiting phase-locking alters recognition in task-dependent ways [43], all consistent with temporal mechanisms dominating while fiber loss is spared, and with the frequency dependence expected from neural-fluctuation coding [44]. Different peripheral pathologies also produce distinguishable cortical signatures [45], and the premise our framework operationalizes, that neural degeneration invisible to the audiogram drives intelligibility deficits, is established both behaviorally [46] and histopathologically [47]. At the level of the test, Beechey [48] showed that intelligibility tests capture only part of communication; Experiment 4 identifies a complementary narrowing on the scoring side, and here, unlike in patients, the mechanism is known, so the loss can be measured. That phoneme-level scoring outperforms word-level scoring for classifying hearing-loss severity [49, 50], and that token confusions already localize impaired channels and distinguish strategies in cochlear implants [24, 51], confirms in patients what we quantify in simulation: a less aggregated score recovers power the word score loses, which at the word level is further diluted by lexical context that lets top-down knowledge repair missing phonemes [52].

### Clinical implications

The limit exposed by Experiment 4 is in the analysis, not the material: larger patient cohorts scored the same way would inherit the same blindness. The full confusion matrix is impractical to collect clinically, but it is needed only during discovery: characterizing the complete structure under known mechanisms identifies the few directional contrasts an efficient test ultimately measures. This motivates a shift from phonetically balanced to phonetically efficient assessment, following the logic of optimal experimental design [53, 54] rather than phonetic balance [55]. This shift is specific to mechanism identification in auditory nerve disorders: for regular sensorineural hearing loss, where broadened cochlear filters degrade audibility while largely respecting temporal cues, phonetic balance remains essential for clinical speech audiometry. The convergence is not an artifact of the word list: the FraMatrix is already highly balanced, so its collapse of the temporal mechanisms shows the limit lies at the word level itself. A single test built around the discriminative contrasts, scored by those contrasts rather than as one percentage, suffices to separate the temporal mechanisms; designing and validating such a list is future work.

A practical consequence for rehabilitation follows: processing that relies on temporal integration preserves benefit for fiber loss but degrades the temporal mechanisms severely (Table 3), so mechanism identification is a prerequisite for safe, targeted intervention [56]. In practice the two often co-occur: the common mixed presentation is an auditory nerve disorder from one of the mechanisms modeled here together with regular sensorineural hearing loss from outer hair cell failure, which disrupts audibility but not temporal cues, so identifying the neural component matters even when peripheral loss is also present.

### Binaural processing

The reasoning that follows is expected on mechanistic grounds but is not tested in the present work. Interaural time differences are resolved on the order of tens of microseconds, requiring bilaterally synchronized inputs to brainstem coincidence detectors. The same neural desynchronization that scrambles monaural timing also disrupts this bilateral alignment, so binaural unmasking, spatial release from masking, and interaural integration need not be restored even when peripheral integrity appears preserved. Monaural audibility can survive while the binaural advantage collapses, because the two draw on different temporal tolerances, a dissociation that depends on which aspect of neural encoding is disrupted, not on whether the synapse itself is intact.

## Limitations

We modeled high-SR fibers only. The fiber-loss perturbation therefore represents frequency-channel elimination (analogous to dead regions) rather than the selective low/medium-SR synaptopathy of Kujawa and Liberman [18, 19]; mixed-SR simulations would be needed to capture SR-specific loss and to test whether fiber loss and truncation remain separable under a more realistic population model. The neurogram perturbations manipulate neural representations directly without modeling the biophysics that produce them; bridging to parameter-based models [26, 32] would enable predictions from histological data. Formant tracking applied to non-resonant sounds indexes broadband spectral dynamics rather than true resonances, consistent with the weaker acoustic-neural correspondence we observe for those categories. The ASR classifier is a feedforward ideal observer that sets an upper bound on recoverable information without capturing central integration, top-down prediction, or perceptual learning. The contrast-based recovery is a proof of concept: the true inter-listener variability, the handling of mixed mechanisms, and the efficient contrast list remain to be validated in listeners. Modeling high-SR fibers only may also be less limiting for supra-threshold coding than it first appears: high-SR fibers continue to transmit cross-frequency profiles of neural fluctuation even when their average rate saturates, so rate saturation does not imply information saturation [44]. On this view the distinctive contribution of low and medium-SR fibers may lie partly in efferent gain control rather than in direct afferent coding, an axis our model does not represent. Relatedly, the truncation perturbation models a loss of representational dynamic range, not the physiological synaptic adaptation that can itself sharpen fluctuation contrast, and the two should not be conflated.

Two validation paths exist: collecting confusion matrices from ANSD patients with characterized pathophysiology, and using the model prospectively to design mechanism-specific test batteries. The harder step is inversion: multiple mechanisms can produce overlapping patterns and patients may present mixed pathology, so confident assignment will require complementary tests that fail differently under different mechanisms.

### Conclusions

Clinical speech scores measure how much is understood but discard the structure that reveals why errors occur. Aggregate scoring removes most of the mechanism-discriminative information the confusion matrix preserves, yet a few targeted contrasts recover it, so the diagnostic gap is one of analysis, not of data collection. Reading the structure of speech errors rather than their aggregate is a step toward mechanism-stratified assessment. More broadly, the framework provides a quantitative way to evaluate whether behavioral summaries preserve the mechanistic information they are intended to represent.

## 4 Materials and Methods

### Auditory nerve modeling

Auditory nerve responses were generated using the Zilany et al. model [32]: 150 logarithmically-spaced characteristic frequencies (125 Hz–10 kHz), high spontaneous rate fibers (100 spikes/s), 65 dB SPL stimulus level. Temporal resolution: 2 ms (500 Hz sampling).

### Speech stimuli

Speech materials were drawn from the TIMIT corpus [31], using the standard train/test partition: 462 training speakers, 168 test speakers, no overlap between sets. Phonemes were grouped into 9 categories (Vowel, Nasal, Liquid, Glide, Fricative, Stop, Affricate, Flap, Silence). For noise-trained models, additive noise from the WHAM! corpus [57] was mixed with speech at SNRs uniformly sampled from 0–20 dB.

### Neural perturbations

Four types modeled ANSD mechanisms: *Uniform jitter*: delay *δ*_*t*_ ∼ *U* (3, 10) ms applied identically across all frequency channels (preserving cross-frequency timing as reference condition). *Scattered jitter*: independent delays *δ*_*t*,*f*_ ∼ *U* (3, 10) ms per channel (destroying cross-frequency synchrony). *Fiber loss*: 1–4 of 150 frequency channels zeroed. *Truncation*: amplitude limited at ratio *α* ∼ *U* (0.3, 0.7) of maximum. See SI Appendix for mathematical definitions and physiological rationale.

#### Experiment 1: optimal transport

We asked whether acoustic similarity structure is preserved in neural representations. Because acoustic features (formant frequencies in Hz) and neural responses (firing rates in spikes/s) live in different spaces and cannot be compared directly, we used the Gromov-Wasserstein (GW) distance [58, 33], which compares the *relationships* within each space rather than raw values: zero means acoustic similarity structure is perfectly preserved in the neural code, larger values mean greater distortion. For each phoneme category and perturbation type, we computed the GW distance between two similarity matrices, one of pairwise acoustic distances (from formant trajectories) and one of pairwise neural distances (from neurograms), yielding a single number per category and perturbation. Formant trajectories (F1, F2, F3) were estimated with the deep-learning, probabilistic heat-map tracker of Shrem et al. [59], applied uniformly to all phoneme segments; for non-resonant categories (stops, affricates, fricatives) these trajectories index broadband spectral dynamics and formant transitions rather than vocal-tract resonances (see Limitations). The formal GW definition, the entropic-regularization parameters, and implementation details are given in the SI Appendix.

#### Experiment 2: speech recognition and misclassification analysis

Following [36], we used a two-stage hierarchical automatic speech recognition system: Stage 1 extracts acoustic-phonetic features from brief windows; Stage 2 integrates predictions across longer temporal context. Stage 1 (456K parameters): three-layer unidirectional GRU (128 units), multi-head attention (4 heads), dense layers with dropout (0.3); input: 50-frame windows (100 ms). Stage 2 (406K parameters): three-layer bidirectional GRU (192, 192, 128 units), multi-head attention (6 heads), dense layers with dropout (0.4); input: 305-frame sequences (610 ms context). Output: 9 phoneme categories. Training used the Adam optimizer, focal loss (*γ* = 2), and early stopping; full training procedures and hyperparameters are in the SI Appendix.

To identify which neural features drive the misclassifications, we adapted a reverse-correlation approach [38]: averaging the neurograms that lead the ASR to predict a given category recovers the characteristic neural pattern, or “template”, for that prediction. Because the ASR outputs a probability per category, we used a confidence-weighted average for each true-predicted phoneme pairing,

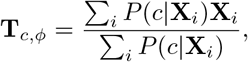

where the sum runs over all tokens of phoneme *ϕ* and *P* (*c* | **X**_*i*_) is the ASR probability for category *c* given neurogram **X**_*i*_, so that **T**_*c,ϕ*_ is the typical pattern of *ϕ* that makes the classifier predict *c*. Difference maps between templates from distinct population and classification-outcome conditions quantify how ANSD perturbations redistribute energy across time and frequency. Analysis focused on stop /t/ and vowels /aa/, /ih/, /iy/; additional consonants are in the SI Appendix.

#### Experiment 3: noise robustness

The four trained models were evaluated in silence and in noise. Silence-trained and noise-trained variants were compared, and accuracy was measured across signal-to-noise ratios (from +15 to − 5 dB) using additive WHAM! noise [57]. To isolate individual mechanisms, the same models were additionally tested on each perturbation type applied alone, rather than on the mixed-perturbation set used for training. Signal-to-noise-dependent and mechanism-isolation results are reported in the SI (Tables S11 and S13).

#### Experiment 4: clinical scoring analysis

The Stage 2 ANSD-Silence confusion matrices were used both as fixed mechanism signatures and as probabilistic models of virtual listeners, from which synthetic responses to the held-out TIMIT sentences were generated with the mechanism known by construction. We then summarized these responses at progressively coarser levels and measured how much mechanism information each level preserves. Full numerical results and parameter settings are in the SI (Clinical Scoring Simulation section).

##### Confusion-matrix representations and scoring

Two quantities were read deterministically from each mechanism’s matrix: the overall score (mean of the diagonal) and error routing (the fraction of misclassified consonant tokens assigned to vowels, other consonants, or silence); between-mechanism differences are pointwise differences between row-normalized matrices. To simulate clinical scoring, each matrix was treated as a categorical response model, drawing the perceived category for a presented phoneme *ϕ* from row **C**_*ϕ*,:_; deterministic scoring takes the maximum-probability response, and we report stochastic scoring (responses sampled) unless otherwise noted. Word and sentence scores were computed under five scoring rules of increasing tolerance (strict, two error-tolerant, two sigmoidal), applied identically to all mechanisms and bracketing plausible clinical rules, in a simulated FraMatrix-style closed-set matrix sentence test (five word positions, approximately ten alternatives each; 2,000 trials per mechanism; SI Table S14). Two robustness checks are reported in the SI: a cross-transfer variant using cross-population matrices (SI Table S15) and an open-set VCV variant (SI Table S17).

##### Mutual information

Treating mechanism identity *M* as uniform over the four types (*H*(*M*) = 2 bits), we estimated *I*(*M* ; *X*) at four levels of aggregation, the full confusion matrix, the per-category accuracy vector, the phoneme-level score, and the word-level score, from 5,000 stochastic test sessions per mechanism (silence excluded, so the matrix is 8 by 8). *I*(*M* ; *X*) retained 78%, 49%, 28%, and 8% of *H*(*M*) across levels (SI Table S16). The absolute percentages depend on estimation parameters, but the monotonic ordering and the conclusion that aggregation discards most mechanism information are robust to these choices. As a binning-free check, a four-class nearest-centroid classifier under five-fold cross-validation reached 76.7% from the full matrix versus 53.1% from the aggregate score (25% baseline), giving *I*(*M* ; *X*) ≥ 0.85 bits via Fano’s inequality [60].

##### Contrast-based classification

Separability of truncation and jitter was tested with a nearest-centroid classifier [61, 62] on the scalar aggregate score versus the vectorized full matrix, against a 500-permutation null (Fig. 5C).

To test how few entries suffice to separate the three temporal mechanisms, we then simulated listeners with three sources of variability rather than one matrix per mechanism: inter-listener variation (confusion-matrix rows drawn from Dirichlet distributions centered on the mechanism matrix [25]), a variable severity level warping the matrix between near-diagonal and reduced-diagonal forms, and a variable number of responses per category. A nearest-centroid rule under five-fold cross-validation, with both the discriminative contrasts (the *K* highest-variance off-diagonal cells across the three centroids) and the centroids estimated on training folds only, classified the mechanisms; accuracy was averaged over folds and seeds, a label-permutation control confirmed chance performance under permuted identities, and mixed-mechanism listeners (convex combinations of two matrices) were classified with the same rule. Structural distance between mechanism matrices was also quantified with the Gromov-Wasserstein distance [58, 33, 63], giving the 6.7 *×* structural-versus-aggregate separability gain for truncation versus jitter (SI Table S18). The variability model and the handling of mixtures are not estimated from data here; a fuller generative and optimal-design treatment is developed separately.

### Statistical testing

Significance of category-specific accuracy differences was assessed using McNemar’s test on paired token-level predictions within each phoneme category. For each comparison (e.g., Healthy-Silence vs ANSD-Silence on stop consonants), a 2 *×* 2 contingency table categorised tokens by whether each model classified them correctly. Bootstrap 95% confidence intervals were computed using 10,000 resamples with replacement (percentile method), preserving token pairing so the same tokens appear in both model predictions within each resample. All reported comparisons were significant at *p <* 0.001 unless otherwise noted. Complete statistical results are provided in SI Tables S6–S10.

## Data Availability

TIMIT corpus available from Linguistic Data Consortium (LDC93S1). WHAM! noise corpus at http://wham.whisper.ai/. Analysis code and trained models available at https://github.com/mcampi111/SimsACues_ASR.

## Contributions

M.C.: Conceptualization, data curation, formal analysis, investigation, methodology, software, writing, original draft, writing, review & editing.

E.P.: Conceptualization, formal analysis, investigation, writing, original draft, writing, review & editing.

G.G.: Formal analysis, investigation, writing, review & editing.

P.A.: Conceptualization, investigation, supervision, funding acquisition, writing, review & editing.

C.G.: Conceptualization, formal analysis, methodology, writing, review & editing.

## Acknowledgments

This work was supported by a grant from Fondation Pour l’Audition (FPA) to the Institut de l’Audition (FPAIDA09 to Hung Thai-Van and FPAIDA10 to Paul Avan and the Ceriah) and a French government grant managed by the Agence Nationale de la Recherche under the France 2030 program, reference ANR-23-IAHU-0003. CG was supported by the Pasteur-Roux-Cantarini Fellowship from Institut Pasteur.

MC thanks Prof. Birger Kollmeier for guidance, Dr. Léo Varnet and Dr. Emmanuel Ponsot for template analysis feedback, and Prof. Romain Serizel for ASR suggestions.

