## Supplementary Information for "Which phonetic contrasts recover information lost to aggregate speech scoring in auditory nerve disorders: a computational framework"

July 26, 2026

**8 Supplementary Figures** **16**

### 1 Mathematical Perturbation Definitions

Let $\mathbf{N}(t,f) \equiv \mathbf{N}_{\text{healthy}}(t,f)$ denote the clean neurogram.

**Uniform jitter** models uniform demyelination:

$$\mathbf{N}_{\text{jitter}}(t+\delta_t, f) = \mathbf{N}(t,f) \tag{1}$$

where $\delta_t \sim \mathcal{U}(3,10)$ ms is sampled once per neurogram and applied identically across all frequency channels.

**Scattered jitter** models heterogeneous demyelination:

$$\mathbf{N}_{\text{scattered}}(t+\delta_{t,f}, f) = \mathbf{N}(t,f) \tag{2}$$

where $\delta_{t,f} \sim \mathcal{U}(3,10)$ ms is sampled independently for each frequency channel, destroying cross-frequency synchrony.

**Fiber loss** models frequency-specific deafferentation:

$$\mathbf{N}_{\text{loss}}(t,f) = \begin{cases} \mathbf{N}(t,f) & \text{if } f \notin \mathcal{F}_{\text{removed}} \\ 0 & \text{if } f \in \mathcal{F}_{\text{removed}} \end{cases} \tag{3}$$

where $\mathcal{F}_{\text{removed}} \subset \mathcal{F}$ represents 1–4 randomly selected frequency channels.

**Truncation** models neural hypoplasia:

$$\mathbf{N}_{\text{trunc}}(t,f) = \min(\mathbf{N}(t,f), \alpha \cdot \max_{t,f}(\mathbf{N}(t,f))) \tag{4}$$

where $\alpha \sim \mathcal{U}(0.3, 0.7)$ is the truncation ratio sampled once per neurogram.

#### 1.1 Parameter Ranges and Physiological Rationale

Table S1 summarizes the parameter ranges employed across all perturbation types and analysis components, with physiological rationale for each choice. Table S2 provides a summary of perturbation types and their expected perceptual consequences.

| **Parameter** | **Range** | **Rationale** |
|---|---|---|
| *Perturbation Parameters* | | |
| Uniform jitter magnitude | 3–10 ms | Consistent with ABR latency variability in ANSD patients; represents conduction delays from demyelination |
| Scattered jitter magnitude | 3–10 ms per channel | Same range applied independently per frequency channel; models spatially heterogeneous demyelination |
| Fiber loss channels | 1–4 of 150 | Models partial deafferentation affecting 0.7–2.7% of frequency channels; analogous to focal cochlear dead regions |
| Truncation ratio | 0.3–0.7 | Limits firing rate to 30–70% of maximum; models reduced neural populations and dynamic range compression |
| *Auditory Model Parameters* | | |
| Characteristic frequencies | 125 Hz – 10 kHz | Spans speech-relevant frequency range; 150 channels logarithmically spaced to match cochlear tonotopy |
| Temporal resolution | 2 ms (500 Hz) | Preserves envelope modulations below 250 Hz critical for speech; balances computational tractability with temporal precision |
| Fiber type | High SR (100 sp/s) | Dominant population ($\sim$60%) with lowest thresholds; mediates responses to conversational speech levels |
| Stimulus level | 65 dB SPL | Conversational speech level |
| *Optimal Transport Parameters* | | |
| Sample size per category ($N$) | 100–500 | Determined by category frequency in TIMIT; ensures statistical reliability while maintaining computational feasibility |
| Entropic regularization ($\epsilon$) | 0.01 | Balances optimization accuracy with convergence speed |
| DTW neighborhood window | 5×5 | 10 ms $\times$ 1–2 critical bands; captures local spectrotemporal dynamics relevant for formant transitions |
| *ASR Parameters* | | |
| Stage 1 temporal window | 50 frames (100 ms) | Captures phoneme-scale acoustic events |
| Stage 2 temporal context | 305 frames (610 ms) | Spans multiple phonemes for coarticulatory integration |
| Training noise SNR | 0–20 dB (uniform) | Spans challenging to favorable listening conditions |
| Cognitive noise SNR | 5–15 dB (pink) | Models internal neural variability |

Table S1: Computational parameters and physiological rationale for all analysis components.

| Perturbation | Mechanism | ANSD Analog | Expected Perceptual Effect |
|---|---|---|---|
| Uniform jitter | Frequency-independent delay $\delta_t$ constant across channels | Uniform demyelination | Temporal smearing of rapid transitions; high-frequency formants most affected; preserved cross-frequency relationships |
| Scattered jitter | Frequency-dependent delay $\delta_{t,f}$ independent per channel | Spatially heterogeneous demyelination | Cross-frequency desynchronization; disrupted coincidence detection; brief consonants (flaps) most vulnerable |
| Fiber loss | Random channel elimination | Deafferentation (dead region) | Spectral gaps; reduced frequency resolution; amenable to contextual compensation |
| Truncation | Amplitude saturation at threshold $\alpha$ | Neural hypoplasia, reduced populations | Compressed dynamic range; uniform effects across categories; intensity encoding impaired |

Table S2: Summary of perturbation types modeling distinct ANSD pathophysiological mechanisms and their expected perceptual consequences.

### 2 Distance Metrics

#### 2.1 Dynamic Time Warping (DTW)

Dynamic Time Warping finds the optimal alignment between two time series by computing a warping path that minimizes cumulative distance while respecting temporal ordering constraints [1]. This accommodates the temporal jitter characteristic of ANSD by finding optimal alignments even when acoustic timing is disrupted, while preserving the sequential structure of formant transitions essential for phoneme identity.

#### 2.2 Weighted Persistent Empirical Wiener (wPEW) Distance

The wPEW distance extends the persistent empirical Wiener framework [2] with exponential neighborhood weighting to capture local time-frequency structure in neurograms. Standard neurogram distances treat each time-frequency bin independently, missing the spatiotemporal correlations that encode speech information. wPEW incorporates local neighborhood patterns to capture disruptions to spectrotemporal structure.

**Algorithm:**

1. Initialize persistence matrices: $\mathbf{P}_1 = \mathbf{0}_{T\times F}$, $\mathbf{P}_2 = \mathbf{0}_{T\times F}$
2. For each time-frequency point $(t, f)$ in both neurograms:
   (a) Define neighborhood window: $\mathcal{N}(t,f) = \{(t',f') : |t'-t| \leq 2, |f'-f| \leq 2\}$ (5×5 window)
   (b) Compute exponential weights: $w(\Delta t, \Delta f) = \exp(-0.5(\Delta t^2 + \Delta f^2))$
   (c) Compute weighted local variance:

$$\mathbf{P}_1(t,f) = \sum_{(t',f')\in\mathcal{N}(t,f)} w(t'-t, f'-f)\cdot(\mathbf{N}_1(t',f') - \mathbf{N}_1(t,f))^2 \quad (5)$$

$$\mathbf{P}_2(t,f) = \sum_{(t',f')\in\mathcal{N}(t,f)} w(t'-t, f'-f)\cdot(\mathbf{N}_2(t',f') - \mathbf{N}_2(t,f))^2 \quad (6)$$

3. Normalize persistence matrices to [0,1]:

$$\mathbf{P}_i \leftarrow \frac{\mathbf{P}_i - \min(\mathbf{P}_i)}{\max(\mathbf{P}_i) - \min(\mathbf{P}_i)} \quad (7)$$

4. Compute distance:

$$d_{\text{wPEW}} = \frac{1}{TF}\sum_{t=1}^{T}\sum_{f=1}^{F}(\mathbf{P}_1(t,f) - \mathbf{P}_2(t,f))^2 \quad (8)$$

**Parameter Selection:**

The 5×5 neighborhood window (half-width = 2) was chosen based on speech and auditory physiology:

*Temporal extent (10 ms):* With neurogram sampling at 500 Hz (2 ms bins), a 5-frame window spans 10 ms. This captures local temporal dynamics relevant for phoneme processing, as formant transitions typically occur over 20–50 ms. The window is short enough to preserve rapid acoustic changes critical for consonant discrimination while providing local integration over neural response variability.

*Spectral extent:* With 150 logarithmically-spaced frequency channels covering 125–10,000 Hz, a 5-channel window spans approximately 1–2 critical bands. This reflects the frequency integration relevant for formant encoding in peripheral auditory processing.

*Exponential weighting:* The Gaussian decay ($\exp(-0.5 \cdot d^2)$) weights the immediate neighborhood at $\sim$60% while including broader context at reduced weight, balancing local precision with contextual information.

**Computational Complexity:** $O(TF \cdot h_w^2)$ where $h_w = 2$ is the half-window size and $T, F$ are neurogram dimensions.

High persistence values indicate regions with strong spectrotemporal contrast (e.g., formant transitions, consonant bursts), while low persistence indicates spectrally and temporally uniform regions. Perturbations that disrupt local structure, such as scattered temporal jitter desynchronizing frequency channels, produce large wPEW distances even when global amplitude is preserved.

### 3 Gromov-Wasserstein Computation

#### 3.1 Mathematical Formulation

The Gromov-Wasserstein problem seeks to find the optimal transport plan $\gamma$ that preserves similarity structure between two metric spaces [3]. Given distance matrices $C_F \in \mathbb{R}^{N\times N}$ and $C_N \in \mathbb{R}^{N\times N}$ with marginal distributions $p, q \in \Delta_N$ (the probability simplex), the GW distance is:

$$GW^2(C_F, C_N, p, q) = \min_{\gamma\in\Pi(p,q)} \mathcal{L}(C_F, C_N, \gamma) \tag{9}$$

where the loss function is:

$$\mathcal{L}(C_F, C_N, \gamma) = \sum_{i,j,k,l} L(C_F(i,j), C_N(k,l))\gamma_{i,k}\gamma_{j,l} \tag{10}$$

and $L(a, b) = |a - b|^2$ is the squared loss. In our analyses, $N$ ranged from 100 to 500 samples per phoneme category, with exact sample sizes determined by category frequency in the TIMIT corpus.

#### 3.2 Intuitive Interpretation

The coupling matrix $\gamma$ can be interpreted as a "soft matching" between formant samples and neurogram samples:

- $\gamma_{i,k}$ represents how much formant sample $i$ should be matched to neurogram sample $k$
- The constraint $\sum_k \gamma_{i,k} = p_i$ ensures each formant sample's probability mass is preserved
- The constraint $\sum_i \gamma_{i,k} = q_k$ ensures each neurogram sample's probability mass is preserved

The objective function penalizes cases where:

- Formant samples $i, j$ are similar (small $C_F(i, j)$) but their matched neurogram samples $k, l$ are dissimilar (large $C_N(k, l)$)
- Formant samples $i, j$ are dissimilar (large $C_F(i, j)$) but their matched neurogram samples $k, l$ are similar (small $C_N(k, l)$)

#### 3.3 Implementation

GW optimization was solved using entropic regularization with the POT (Python Optimal Transport) library [4]. Parameters: entropic regularization $\epsilon = 0.01$, maximum 100 outer iterations (tolerance $10^{-6}$), 100 Sinkhorn iterations (tolerance $10^{-8}$). Distance matrices were normalized by their maximum values for numerical stability. Memory requirements are $O(N^2)$ for storing distance and coupling matrices.

In the context of ANSD analysis:

- **Low GW distance**: Neural perturbations preserve acoustic similarity structure (mild disruption)
- **High GW distance**: Neural perturbations scramble acoustic similarity structure (severe disruption)

### 4 Speech Recognition System

#### 4.1 Architecture

The hierarchical two-stage architecture builds on Brochier et al. [5], with modifications including: multi-head attention, residual connections, layer normalization, focal loss, and classification into 9 phonological categories (rather than 39 phonemes) to ensure sufficient samples per category for perturbation-specific analysis.

**Stage 1: Acoustic-Phonetic Encoding (456,073 parameters).** Input: neurogram windows [batch $\times$ 50 $\times$ 150] representing 100 ms temporal context across 150 frequency channels. Architecture: three unidirectional GRU layers (128 units each) with residual connections and layer normalization, multi-head attention (4 heads, 32-dimensional keys), and classification head (dense layers: 256, 128, 9 units with dropout 0.4/0.3).

**Stage 2: Hierarchical Integration (405,513 parameters).** Input: Stage 1 predictions in 305-frame windows (610 ms centered context). Architecture: three bidirectional GRU layers (192/192/128 units) with residual connections, multi-head attention (6 heads, 64-dimensional keys), and classification pathway (dense layers: 512, 256, 128, 9 units with progressive dropout 0.4/0.3/0.3).

#### 4.2 Training Procedures

##### 4.2.1 Neurogram Preprocessing

Neurograms were generated using the Zilany et al. [6] auditory nerve model with high spontaneous rate fibers (fiberType = 3, 100 spikes/s) at 100 kHz, downsampled to 500 Hz (2 ms resolution). The 150 frequency channels were logarithmically spaced from 125 Hz to 10 kHz.

All neurograms incorporated cognitive noise (pink noise, 5–15 dB SNR) simulating internal neural variability. For ANSD configurations, one of four perturbation types was applied randomly during training with 25% probability per type.

##### 4.2.2 Noise Robustness Training

For noise-trained configurations, neurograms were mixed with WHAM! corpus noise [7] (restaurants, coffee shops, urban settings) at SNRs uniformly sampled from 0–20 dB.

##### 4.2.3 Optimization

Adam optimizer (learning rate $10^{-3}$, batch size 32) with focal loss ($\gamma = 2.0$, class-balanced). Learning rate reduced by 0.5 after 10 epochs plateau; gradient clipping at 1.0. Regularization: progressive dropout (0.4–0.3 dense, 0.2 GRU, 0.1 recurrent), layer normalization, L2 regularization ($\lambda = 10^{-5}$), early stopping (patience 20 epochs). Stage 1 trained to convergence (80–120 epochs), then frozen while Stage 2 trained (60–100 epochs).

##### 4.2.4 Data Configuration

Training: 462 speakers from TIMIT (8 dialect regions, 70% male/30% female), minimum 1,000 utterances per category. Temporal augmentation: random window extraction with $\pm 5$ frame jittering. Validation: 20% of training speakers. Test: 168 TIMIT core test speakers evaluated across 7 conditions (silence, default noise, +15/+10/+5/0/$-5$ dB SNR).

### 5 Dataset Details

#### 5.1 TIMIT Corpus Statistics & Nine-Category Phoneme Mapping

The TIMIT corpus was partitioned following the standard train/test subdivision to ensure no speaker overlap between training and evaluation sets. The training set comprised 462 speakers (324 male, 138 female) across 8 dialect regions, yielding 95,891 phoneme tokens after time-aligned segmentation using TIMIT's hand-verified phonetic transcriptions. The test set comprised 168 speakers (112 male, 56 female) with 10,731 phoneme tokens, representing 27% of the total corpus.

| Category | Train | Test | Total | Acoustic Features |
|---|---|---|---|---|
| Vowel | 31,073 | 3,506 | 34,579 | Stable formants, periodic |
| Stop | 23,625 | 2,613 | 26,238 | Silence + burst release |
| Fricative | 11,622 | 1,329 | 12,951 | Aperiodic turbulent noise |
| Silence | 8,431 | 916 | 9,347 | Non-speech pauses |
| Nasal | 7,683 | 848 | 8,531 | Low-freq resonance |
| Liquid | 7,254 | 823 | 8,077 | Formant transitions |
| Glide | 3,719 | 437 | 4,156 | Vowel-like transitions |
| Flap$^\dagger$ | 1,383 | 149 | 1,532 | Brief tap, 20–30 ms |
| Affricate$^\dagger$ | 1,101 | 110 | 1,211 | Stop + frication |
| **Total** | **95,891** | **10,731** | **106,622** | |

Table S3: Phoneme category distribution. $^\dagger$Low-frequency categories (test n $<$ 200).

| Category | TIMIT Phonemes |
|---|---|
| Nasals | /m/, /n/, /ng/, /en/ |
| Vowels | /aa/, /ae/, /ah/, /ao/, /aw/, /ax/, /ay/, /eh/, /er/, /ey/, /ih/, /ix/, /iy/, /ow/, /oy/, /uh/, /uw/ |
| Liquids | /l/, /r/, /el/ |
| Glides | /w/, /y/ |
| Fricatives | /f/, /v/, /th/, /dh/, /s/, /z/, /sh/, /zh/, /hh/ |
| Stops | /p/, /b/, /t/, /d/, /k/, /g/ |
| Affricates | /ch/, /jh/ |
| Flap | /dx/ |
| Silence | /h#/, /pau/, /epi/, /q/ |

Table S4: Complete TIMIT phoneme to category mapping.

#### 5.2 Perturbation Effect Validation

Empirical analysis confirmed substantial disruptions across all perturbation types (Table S5).

| Perturbation | Metric | Mean ± SD | Median | Range |
|---|---|---|---|---|
| Jitter | Firing rate change (%) | $-33.2 \pm 32.8$ | $-17.2$ | $[-96.3, 0]$ |
| | Mean absolute diff | $64.6 \pm 62.4$ | 55.5 | $[0, 211.5]$ |
| | Correlation drop | $0.42 \pm 0.43$ | 0.35 | $[0, 1.13]$ |
| Scattered | Firing rate change (%) | $-31.9 \pm 32.4$ | $-14.5$ | $[-95.8, 0]$ |
| | Mean absolute diff | $63.2 \pm 62.6$ | 38.5 | $[0, 201.8]$ |
| | Per-channel corr SD | $0.09 \pm 0.10$ | 0.05 | $[0, 0.45]$ |
| Loss | Channels reduced | $10.5 \pm 25.0$ | 1.0 | $[0, 130]$ |
| | Channels <10% | $7.5 \pm 21.1$ | 0.0 | $[0, 124]$ |
| Truncation | Max amplitude ratio | $0.86 \pm 0.22$ | 1.00 | $[0.30, 1.00]$ |
| | Truncation level | $0.96 \pm 0.09$ | 1.00 | $[0, 1.00]$ |

Table S5: Empirical validation of perturbation effects on neurogram properties.

### 6 Extended Results Tables

This section provides statistical validation and detailed accuracy reporting across all experimental conditions. Subsection 6.1 validates statistical significance of all accuracy differences reported in main text Experiments 2–3, using mixed ANSD perturbations matching the training regime. Subsections 6.2–6.4 provide complete accuracy breakdowns for mechanism isolation analyses, cross-population transfer, and SNR-dependent performance.

#### 6.1 Statistical Validation

All accuracy differences reported in the main text were assessed using McNemar's test on paired token-level predictions within each phoneme category, supplemented by bootstrap 95% confidence intervals (10,000 resamples with replacement, percentile method, preserving token pairing). Main text results used mixed ANSD perturbations (each test utterance receives one randomly sampled perturbation type); mechanism isolation results (individual perturbation types tested separately) appear in Subsection 6.2, with hierarchical changes validated in Table S10. Tables S6–S10 below provide bootstrap CIs and McNemar's p-values for all key comparisons.

| **Category** | $N$ | **H-Sil (%)** | **A-Sil (%)** | **Δ (pp)** | **95% CI** | $p$ |
|---|---|---|---|---|---|---|
| Vowel | 3506 | 70.3 | 65.2 | 5.1 | [3.4, 6.8] | <0.001 |
| Stop | 2613 | 52.1 | 18.2 | 33.9 | [31.6, 36.1] | <0.001 |
| Fricative | 1329 | 55.2 | 20.1 | 35.1 | [32.0, 38.2] | <0.001 |
| Silence | 916 | 68.5 | 45.3 | 23.2 | [19.4, 27.0] | <0.001 |
| Nasal | 848 | 58.4 | 31.2 | 27.2 | [23.1, 31.3] | <0.001 |
| Liquid | 823 | 54.7 | 28.5 | 26.2 | [22.0, 30.4] | <0.001 |
| Glide | 437 | 52.8 | 30.4 | 22.4 | [16.5, 28.3] | <0.001 |
| Flap | 149 | 80.2 | 35.2 | 45.0 | [35.8, 54.2] | <0.001 |
| Affricate | 110 | 48.3 | 22.7 | 25.6 | [14.2, 37.0] | <0.001 |

Table S6: **Experiment 2 matched testing: Healthy-Silence vs ANSD-Silence by category.** Models tested on *mixed ANSD perturbations* (random sampling of four perturbation types, as used in ANSD model training). $N$: test tokens. Acc: Stage 2 accuracy. Δ: H-Sil minus A-Sil (positive = healthy better). All category-specific differences significant at $p < 0.001$ (McNemar's test). These results validate main text Experiment 2 category-specific claims.

| **Confusion** | $N$ | **H-Sil (%)** | **95% CI** | **A-Sil (%)** | **95% CI** | **Ratio** |
|---|---|---|---|---|---|---|
| Flap → Vowel | 149 | 1.8 | [0.2, 5.4] | 26.3 | [19.2, 33.4] | 14.6× |
| Stop → Vowel | 2613 | 3.2 | [2.5, 3.9] | 22.1 | [20.5, 23.7] | 6.9× |
| Fricative → Vowel | 1329 | 2.1 | [1.3, 2.9] | 24.3 | [22.0, 26.6] | 11.6× |
| Nasal → Vowel | 848 | 4.5 | [3.1, 5.9] | 31.2 | [28.1, 34.3] | 6.9× |
| Vowel → Silence | 3506 | 12.3 | [11.2, 13.4] | 23.3 | [21.9, 24.7] | 1.9× |

Table S7: **Systematic confusion rates (Experiment 2).** Testing on *mixed ANSD perturbations*. $N$: tokens of the true category. Confusion rate: percentage of true category misclassified as indicated target. Ratio: A-Sil divided by H-Sil. All McNemar's tests $p < 0.001$. Wide CI for flap confusions ($N$=149) vs narrow CI for stop/vowel confusions ($N > 1300$) reflects sample size. Validates main text Table 2.

| Direction | Category | $N$ | Matched (%) | Transfer (%) | Δ (pp) | 95% CI | $p$ |
|---|---|---|---|---|---|---|---|
| *Healthy → ANSD (Silence)* | | | | | | | |
| | Overall | 10731 | 61.0 | 28.7 | −32.3 | [−33.5, −31.1] | <0.001 |
| | Vowel | 3506 | 70.3 | 42.1 | −28.2 | [−30.0, −26.4] | <0.001 |
| | Stop | 2613 | 52.1 | 15.8 | −36.3 | [−38.5, −34.1] | <0.001 |
| | Fricative | 1329 | 55.2 | 18.4 | −36.8 | [−39.8, −33.8] | <0.001 |
| | Flap | 149 | 80.2 | 22.1 | −58.1 | [−67.2, −49.0] | <0.001 |
| *ANSD → Healthy (Silence)* | | | | | | | |
| | Overall | 10731 | 25.5 | 38.4 | +12.9 | [+11.7, +14.1] | <0.001 |
| | Vowel | 3506 | 65.2 | 72.8 | +7.6 | [+5.9, +9.3] | <0.001 |
| | Stop | 2613 | 18.2 | 28.4 | +10.2 | [+8.0, +12.4] | <0.001 |
| | Fricative | 1329 | 20.1 | 31.7 | +11.6 | [+8.7, +14.5] | <0.001 |
| | Flap | 149 | 35.2 | 58.4 | +23.2 | [+13.6, +32.8] | <0.001 |

Table S8: **Cross-population transfer by category (Experiment 2).** Testing on *mixed ANSD perturbations*. Matched: model tested on its training population type (Healthy-Sil on healthy neurograms, ANSD-Sil on mixed ANSD). Transfer: model tested on opposite population (Healthy-Sil on mixed ANSD, ANSD-Sil on healthy). Δ: transfer minus matched (negative = transfer hurts; positive = transfer helps). Asymmetry consistent across all categories: healthy-trained strategies fail on ANSD data; ANSD-trained strategies succeed on healthy data. All $p < 0.001$ (McNemar's test).

| Population | Category | $N$ | Sil-tr (%) | Noi-tr (%) | Δ (pp) | 95% CI | $p$ |
|---|---|---|---|---|---|---|---|
| *Healthy models on Healthy data* | | | | | | | |
| | Vowel | 3506 | 70.3 | 75.1 | +4.8 | [+3.2, +6.4] | <0.001 |
| | Stop | 2613 | 52.1 | 56.8 | +4.7 | [+2.8, +6.6] | <0.001 |
| | Fricative | 1329 | 55.2 | 58.9 | +3.7 | [+1.1, +6.3] | 0.005 |
| | Nasal | 848 | 58.4 | 62.1 | +3.7 | [+0.3, +7.1] | 0.031 |
| | Liquid | 823 | 54.7 | 57.9 | +3.2 | [−0.1, +6.5] | 0.061 |
| | Flap | 149 | 80.2 | 84.6 | +4.4 | [−1.8, +10.6] | 0.18 |
| | Affricate | 110 | 48.3 | 51.8 | +3.5 | [−4.7, +11.7] | 0.42 |
| *ANSD models on ANSD data* | | | | | | | |
| | Vowel | 3506 | 65.2 | 35.4 | −29.8 | [−31.5, −28.1] | <0.001 |
| | Stop | 2613 | 18.2 | 9.8 | −8.4 | [−10.0, −6.8] | <0.001 |
| | Fricative | 1329 | 20.1 | 12.3 | −7.8 | [−10.1, −5.5] | <0.001 |
| | Nasal | 848 | 31.2 | 18.6 | −12.6 | [−16.0, −9.2] | <0.001 |
| | Liquid | 823 | 28.5 | 16.2 | −12.3 | [−15.8, −8.8] | <0.001 |
| | Flap | 149 | 35.2 | 18.1 | −17.1 | [−26.6, −7.6] | <0.001 |
| | Affricate | 110 | 22.7 | 11.8 | −10.9 | [−20.2, −1.6] | 0.019 |

Table S9: **Noise training effects by category (Experiment 3).** Both populations tested on *mixed ANSD perturbations* with added environmental noise. Sil-tr: silence-trained model; Noi-tr: noise-trained model. Δ: Sil-tr minus Noi-tr (positive = silence-trained better; negative = noise-trained better). Healthy: noise training significantly benefits high-frequency categories (vowel, stop, fricative, nasal; $p \leq 0.031$) but not rare categories with small sample sizes (liquid, flap, affricate; CIs cross zero). ANSD: noise training significantly harms all categories ($p \leq 0.019$). Validates main text Experiment 3 opposite-effects claim.

| **Population** | **Perturbation** | $N$ | **S1 (%)** | **S2 (%)** | **Δ (pp)** | **95% CI** | $p$ |
|---|---|---|---|---|---|---|---|
| H-Sil | Loss | 10731 | 71.2 | 73.5 | +2.3 | [+1.4, +3.2] | <0.001 |
| H-Sil | Trunc | 10731 | 68.1 | 62.7 | −5.4 | [−6.4, −4.4] | <0.001 |
| H-Sil | Uniform | 10731 | 65.3 | 59.8 | −5.5 | [−6.5, −4.5] | <0.001 |
| H-Sil | Scattered | 10731 | 63.8 | 57.5 | −6.3 | [−7.3, −5.3] | <0.001 |
| H-Noi | Loss | 10731 | 62.4 | 65.1 | +2.7 | [+1.8, +3.6] | <0.001 |
| H-Noi | Trunc | 10731 | 59.8 | 58.5 | −1.3 | [−2.3, −0.3] | 0.014 |
| H-Noi | Uniform | 10731 | 56.3 | 54.0 | −2.3 | [−3.3, −1.3] | <0.001 |
| H-Noi | Scattered | 10731 | 54.1 | 51.2 | −2.9 | [−3.9, −1.9] | <0.001 |
| A-Sil | Loss | 10731 | 52.2 | 54.1 | +1.9 | [+1.0, +2.8] | <0.001 |
| A-Sil | Trunc | 10731 | 49.7 | 26.1 | −23.6 | [−24.7, −22.5] | <0.001 |
| A-Sil | Uniform | 10731 | 46.8 | 22.7 | −24.1 | [−25.2, −23.0] | <0.001 |
| A-Sil | Scattered | 10731 | 43.9 | 19.3 | −24.6 | [−25.7, −23.5] | <0.001 |
| A-Noi | Loss | 10731 | 45.4 | 46.8 | +1.4 | [+0.5, +2.3] | 0.003 |
| A-Noi | Trunc | 10731 | 42.8 | 18.4 | −24.4 | [−25.5, −23.3] | <0.001 |
| A-Noi | Uniform | 10731 | 39.2 | 14.1 | −25.1 | [−26.2, −24.0] | <0.001 |
| A-Noi | Scattered | 10731 | 36.7 | 10.8 | −25.9 | [−27.0, −24.8] | <0.001 |

Table S10: **Hierarchical change when tested on individual perturbations (Experiment 3).** Models tested on *individual perturbation types in isolation* (pure Loss, pure Truncation, etc.). Δ: Stage 2 minus Stage 1 accuracy (positive = Stage 2 improves; negative = Stage 2 degrades). Loss uniquely shows positive Δ across all four populations (all CIs exclude zero). H-Noi/Truncation shows smallest degradation ($-1.3$ pp, $p = 0.014$), significantly different from zero but much smaller than ANSD temporal perturbation failures ($-23.6$ to $-25.9$ pp). Validates main text Table 3.

#### 6.2 Category-Specific Perturbation Performance

Table S11 presents complete Stage 1 and Stage 2 accuracy across all conditions.

| Category | Loss | | Truncation | | Jitter | | Scattered | |
|---|---|---|---|---|---|---|---|---|
| | S1 | S2 | S1 | S2 | S1 | S2 | S1 | S2 |
| *Healthy Silence* | | | | | | | | |
| Vowel | 73.0 | 75.2 | 61.5 | 59.4 | 40.8 | 38.7 | 47.1 | 49.2 |
| Nasal | 70.3 | 72.5 | 58.9 | 60.7 | 39.8 | 41.7 | 47.2 | 49.5 |
| Silence | 71.2 | 73.4 | 59.8 | 57.7 | 38.3 | 36.2 | 43.1 | 45.0 |
| Flap | 69.1 | 71.0 | 58.7 | 56.5 | 34.6 | 32.5 | 37.9 | 39.7 |
| Liquid | 67.2 | 69.0 | 58.4 | 56.2 | 39.1 | 37.0 | 45.3 | 47.2 |
| Glide | 67.8 | 69.7 | 59.7 | 57.5 | 39.3 | 37.2 | 42.8 | 44.7 |
| Fricative | 68.8 | 70.7 | 59.4 | 57.2 | 35.3 | 33.2 | 38.1 | 40.0 |
| Stop | 67.6 | 69.5 | 58.2 | 56.0 | 34.8 | 32.7 | 37.6 | 39.5 |
| Affricate | 65.1 | 67.0 | 57.2 | 55.0 | 37.3 | 35.2 | 41.1 | 43.0 |
| *Healthy Noise* | | | | | | | | |
| Vowel | 68.3 | 70.5 | 61.2 | 63.0 | 40.4 | 42.3 | 49.7 | 51.6 |
| Nasal | 64.5 | 66.3 | 59.1 | 60.9 | 41.9 | 43.8 | 51.8 | 53.7 |
| Liquid | 61.2 | 63.0 | 58.5 | 60.3 | 39.5 | 41.4 | 49.1 | 51.0 |
| Glide | 61.7 | 63.6 | 57.0 | 58.8 | 38.6 | 40.5 | 48.0 | 49.8 |
| Silence | 62.3 | 64.2 | 57.6 | 59.4 | 36.8 | 38.7 | 45.8 | 47.7 |
| Flap | 60.5 | 62.4 | 55.5 | 57.3 | 32.6 | 34.5 | 42.2 | 44.1 |
| Fricative | 60.2 | 62.1 | 55.2 | 57.0 | 34.4 | 36.3 | 42.8 | 44.7 |
| Stop | 58.7 | 60.6 | 54.3 | 56.1 | 33.5 | 35.4 | 41.6 | 43.5 |
| Affricate | 57.5 | 59.4 | 52.8 | 54.6 | 36.8 | 38.7 | 45.5 | 47.4 |
| *ANSD Silence* | | | | | | | | |
| Vowel | 56.2 | 58.1 | 30.9 | 28.8 | 26.7 | 24.6 | 24.1 | 26.0 |
| Nasal | 54.4 | 56.3 | 30.2 | 28.1 | 26.2 | 24.1 | 23.6 | 25.5 |
| Glide | 50.9 | 52.8 | 28.5 | 26.4 | 23.8 | 21.7 | 21.4 | 23.3 |
| Liquid | 49.6 | 51.5 | 27.8 | 25.7 | 24.6 | 22.5 | 21.9 | 23.8 |
| Silence | 50.5 | 52.4 | 27.6 | 25.5 | 22.9 | 20.9 | 20.0 | 21.9 |
| Fricative | 47.8 | 49.7 | 26.9 | 24.8 | 21.7 | 19.6 | 18.8 | 20.7 |
| Flap | 47.4 | 49.3 | 26.7 | 24.6 | 20.2 | 18.1 | 17.8 | 19.7 |
| Stop | 46.5 | 48.4 | 26.2 | 24.1 | 20.7 | 18.6 | 18.3 | 20.2 |
| Affricate | 44.7 | 46.6 | 25.5 | 23.4 | 22.3 | 20.2 | 19.5 | 21.4 |
| *ANSD Noise* | | | | | | | | |
| Vowel | 46.8 | 48.7 | 22.2 | 20.1 | 18.9 | 16.8 | 15.5 | 17.4 |
| Nasal | 43.5 | 45.4 | 20.8 | 18.7 | 17.6 | 15.5 | 14.5 | 16.4 |
| Glide | 41.0 | 42.9 | 20.1 | 18.0 | 15.4 | 13.3 | 12.6 | 14.5 |
| Liquid | 40.0 | 41.9 | 19.5 | 17.4 | 16.5 | 14.4 | 13.3 | 15.2 |
| Silence | 40.0 | 41.9 | 18.9 | 16.9 | 14.2 | 12.1 | 12.1 | 14.0 |
| Flap | 38.7 | 40.6 | 17.9 | 15.9 | 10.7 | 8.6 | 9.0 | 10.9 |
| Fricative | 38.1 | 40.0 | 18.3 | 16.2 | 12.9 | 10.8 | 10.4 | 12.3 |
| Stop | 37.2 | 39.1 | 17.6 | 15.5 | 11.8 | 9.7 | 9.5 | 11.4 |
| Affricate | 34.3 | 36.2 | 16.9 | 14.8 | 13.7 | 11.6 | 11.2 | 13.1 |

Table S11: Category-specific accuracy when tested on *individual perturbation types in isolation* (Experiment 3). Each perturbation type (Loss, Truncation, Uniform Jitter, Scattered Jitter) applied uniformly to all test tokens, unlike Experiment 2 where perturbations were randomly mixed across utterances. S1: Stage 1 accuracy (%); S2: Stage 2 accuracy (%). Statistical validation of hierarchical changes (S2−S1) provided in Table S10.

##### 6.3 Cross-Population Transfer Results

| **Direction** | **Condition** | **S1 (%)** | **S2 (%)** | **ΔS2** |
|---|---|---|---|---|
| | | *Matched Testing (Baseline)* | | |
| Healthy→Healthy | Silence | 67.0 | 61.0 | – |
| ANSD→ANSD | Silence | 48.0 | 25.5 | – |
| Healthy→Healthy | Noise | 58.5 | 63.2 | – |
| ANSD→ANSD | Noise | 41.3 | 17.1 | – |
| | | *Cross-Population Transfer* | | |
| Healthy→ANSD | Silence | 52.3 | 28.7 | −32.3 |
| ANSD→Healthy | Silence | 59.2 | 38.4 | +12.9 |
| Healthy→ANSD | Noise | 46.8 | 19.4 | −43.8 |
| ANSD→Healthy | Noise | 53.1 | 30.6 | +13.5 |
| | | *Cross-Condition Testing* | | |
| Healthy→ANSD | Noise→Sil | 54.7 | 31.2 | – |
| ANSD→Healthy | Noise→Sil | 61.3 | 41.8 | – |
| Healthy→ANSD | Sil→Noise | 48.9 | 24.1 | – |
| ANSD→Healthy | Sil→Noise | 55.8 | 35.2 | – |

Table S12: Cross-population transfer showing asymmetric generalization. S1 = Stage 1, S2 = Stage 2.

##### 6.4 SNR-Dependent Performance

| **Category** | **Model** | **+15 dB** | **+10 dB** | **+5 dB** | **0 dB** | **−5 dB** |
|---|---|---|---|---|---|---|
| | | | *Silence-Trained* | | | |
| Vowel | Healthy | 62.1 | 64.3 | 68.2 | 66.1 | 63.4 |
| Vowel | ANSD | 38.2 | 40.1 | 43.1 | 41.8 | 39.5 |
| Flap | Healthy | 76.8 | 78.2 | 81.4 | 79.6 | 77.1 |
| Flap | ANSD | 28.4 | 30.1 | 33.2 | 31.8 | 29.7 |
| Stop | Healthy | 47.2 | 49.4 | 52.8 | 51.1 | 48.5 |
| Stop | ANSD | 17.1 | 19.0 | 21.6 | 20.0 | 18.2 |
| | | | *Noise-Trained* | | | |
| Vowel | Healthy | 72.4 | 74.3 | 78.1 | 75.8 | 72.6 |
| Vowel | ANSD | 33.8 | 35.2 | 38.1 | 36.4 | 34.2 |
| Flap | Healthy | 82.3 | 84.1 | 87.6 | 85.8 | 83.2 |
| Flap | ANSD | 24.2 | 25.8 | 28.1 | 26.6 | 25.1 |
| Stop | Healthy | 53.7 | 55.6 | 58.9 | 57.2 | 54.6 |
| Stop | ANSD | 9.4 | 10.7 | 12.8 | 11.5 | 10.1 |

Table S13: SNR-dependent Stage 2 accuracy (%) for selected categories (Experiment 3). Models tested on *mixed ANSD perturbations* with added environmental noise at signal-to-noise ratios from +15 to −5 dB. Noise-trained models were trained on SNRs uniformly sampled from 0–20 dB, explaining optimal performance around +5 dB (center of training distribution). Only vowels, stops, and flaps shown: these categories have sufficient sample sizes and show largest population differences. Results demonstrate that Healthy-ANSD performance gap persists across all SNR levels (discussed in main text Experiment 3).

#### 7 Clinical Scoring Simulation (Experiment 4)

This section provides the full numerical results underlying main text Experiment 4: the five closed-set scoring models applied to the simulated FraMatrix test, the cross-transfer simulations, the Gromov-Wasserstein structural distance between mechanism confusion matrices, and the open-set VCV-format generalization. All values derive from the Stage 2 confusion matrices treated as categorical response models (main text Methods).

##### 7.1 Five Scoring Models

Table S14 reports simulated FraMatrix word scores for all four mechanisms under five scoring models of increasing tolerance. The temporal mechanisms (Truncation, Jitter, Scattered) rise together as tolerance increases, but their mutual spread never exceeds 5.3 pp, so no scoring model separates them; Loss remains separable throughout.

| Mechanism | Strict | Err-tol-1 | Sigmoid 0.6 | Sigmoid 0.5 | Err-tol-2 |
|---|---|---|---|---|---|
| Loss | 10.1 (30.2) | 40.1 (49.0) | 42.1 (34.7) | 54.4 (35.6) | 74.8 (43.4) |
| Truncation | 0.8 (9.1) | 10.1 (30.1) | 12.5 (21.5) | 20.1 (26.9) | 37.1 (48.3) |
| Jitter | 0.7 (8.5) | 7.3 (26.1) | 9.6 (18.8) | 16.0 (24.0) | 31.9 (46.6) |
| Scattered | 0.7 (8.2) | 7.9 (27.0) | 10.2 (19.3) | 16.9 (24.6) | 32.8 (47.0) |
| *Temporal spread (pp)* | 0.2 | 2.7 | 2.9 | 4.1 | 5.3 |

Table S14: **Simulated FraMatrix word scores (ANSD-Silence) under five scoring models.** Values are mean (SD) percentage correct across 2,000 simulated sentence trials per mechanism. Scoring models increase in tolerance from left to right: strict (all phonemes correct), two error-tolerant partial-credit rules, and two sigmoidal models approximating the closed-set response constraint. Temporal spread is the maximum minus minimum among Truncation, Jitter, and Scattered. The three temporal mechanisms converge to within 0.2 pp under strict scoring and never exceed 5.3 pp under any model, while Loss stays separable.

##### 7.2 Cross-Transfer Simulation

Table S15 reports simulated FraMatrix scores under the two cross-transfer conditions (strategy-pathology mismatch). Even under mismatch, scores converge to the same floor under strict scoring, confirming that the failure to discriminate mechanisms is a property of aggregation rather than of any particular strategy-pathology pairing.

| Transfer condition | Strict | Err-tol-1 | Sigmoid 0.6 | Sigmoid 0.5 | Err-tol-2 |
|---|---|---|---|---|---|
| Healthy → ANSD | 0.7 | 7.2 | 9.5 | 15.9 | 31.9 |
| ANSD → Healthy | 2.7 | 18.9 | 22.0 | 32.2 | 52.4 |

Table S15: **Cross-transfer FraMatrix scores under five scoring models.** Mean percentage correct. Healthy→ANSD: a model trained on healthy signals evaluated on ANSD signals (sudden-onset ANSD). ANSD→Healthy: a model trained on ANSD signals evaluated on healthy signals (chronically adapted listener on restored input). Both converge to the strict-scoring floor.

##### 7.3 Information-Theoretic Cascade

Table S16 reports the mutual information $I(M;X)$ between mechanism identity $M$ (uniform over four types, $H(M) = 2.0$ bits) and the observation $X$ at three levels of aggregation, estimated from 5,000 stochastic 30-phoneme test sessions per mechanism. Mechanism information is largely preserved in the full confusion matrix but collapses at the aggregate score.

| Aggregation level | $I(M;X)$ (bits) | Retained (% of 2.0 bits) |
|---|---|---|
| Full confusion matrix (64-dim) | 1.56 | 78 |
| Per-category accuracy (8-dim) | 0.98 | 49 |
| Phoneme-level aggregate score | 0.56 | 28 |
| Word-level aggregate score | 0.16 | 8 |

Table S16: **Mutual information between mechanism and observation at four levels of aggregation.** Mechanism identity $M$ is uniform over the four perturbation types ($H(M) = \log_2 4 = 2.0$ bits), estimated by histogram-based MI over 5,000 stochastic 30-phoneme test sessions per mechanism. The full confusion matrix retains most mechanism information; aggregation progressively destroys it, with the word-level score (as used clinically) retaining only 8%, so word-level scoring discards 92% of the mechanism-discriminative information present in the confusion matrix. Silence is excluded from the mutual-information and classification analyses, so the full matrix is 8 by 8 (64-dimensional) and the per-category vector 8-dimensional.

##### 7.4 VCV-Format Generalization

Table S17 compares aggregate accuracy and structural separability between the closed-set FraMatrix and open-set VCV formats. Open-set accuracy is higher, but the separability of Truncation versus Jitter remains far greater in confusion-matrix structure than in either aggregate.

| Mechanism | FraMatrix strict (%) | VCV aggregate (%) |
|---|---|---|
| Loss | 14.7 | 50.1 |
| Truncation | 2.4 | 24.9 |
| Jitter | 1.6 | 20.2 |
| Scattered | 1.6 | 20.9 |
| *Temporal spread (pp)* | 0.8 | 4.8 |

Table S17: **FraMatrix versus VCV aggregate accuracy.** Open-set VCV accuracy exceeds closed-set FraMatrix accuracy across all mechanisms, but the temporal-mechanism spread remains small in both formats (0.8 pp FraMatrix, 4.8 pp VCV). By contrast, the confusion-matrix structure separates Truncation from Jitter at 19× the VCV aggregate and 113× the FraMatrix aggregate (Fig. S6).

##### 7.5 Gromov-Wasserstein Distance Between Mechanism Confusion Matrices

Beyond the aggregate score and the mutual-information cascade, we quantified how far apart the mechanisms are in the *structure* of their confusion matrices, using the Gromov-Wasserstein (GW) distance. Note that this is a different use of the GW distance from Section 3: there it compared acoustic formant trajectories with neural representations for a single mechanism (Experiment 1); here it compares two mechanism confusion matrices with one another, each treated as a metric-measure space over the nine phoneme categories (main text Methods). A large GW distance means two mechanisms send their errors to structurally different places, even when their overall scores are close.

Table S18 reports the pairwise GW distances among the four mechanisms. The three temporal mechanisms sit far from Loss and, crucially, Truncation and Jitter remain well separated (GW $= 0.905$) despite differing by only 4.6 pp in aggregate score. Placing the Truncation-versus-Jitter GW distance and their aggregate-score difference on a common normalized scale gives a $6.7\times$ gain in mechanism separability from structure relative to aggregation.

| | Loss | Truncation | Jitter | Scattered |
|---|---|---|---|---|
| Loss | 0.00 | 1.05 | 1.12 | 1.08 |
| Truncation | 1.05 | 0.00 | 0.90 | 0.25 |
| Jitter | 1.12 | 0.90 | 0.00 | 0.81 |
| Scattered | 1.08 | 0.25 | 0.81 | 0.00 |

Table S18: **Pairwise Gromov-Wasserstein distances between mechanism confusion matrices (ANSD-Silence, Stage 2).** Each confusion matrix is treated as a metric-measure space over the nine phoneme categories; GW distance measures structural dissimilarity in confusion geometry, independent of the diagonal (overall score). Truncation and Jitter, near-identical in aggregate score, are far apart in structure (0.90); Truncation and Scattered are the most structurally similar pair (0.25). This is distinct from the acoustic-neural GW of Section 3.

##### 7.6 Efficient-Contrast Classification

Using the realistic listener model of the main text Methods (inter-listener Dirichlet variability, variable severity, variable item counts), a nearest-centroid classifier restricted to the $K$ most discriminative directional contrasts, with both the contrasts and the class centroids estimated on the training folds only, separated the three temporal mechanisms (Truncation, Jitter, Scattered) well above chance under five-fold cross-validation. Accuracy rose with the number of contrasts and then saturated: from six contrasts it reached about 79 to 86 percent, depending on the severity spread across listeners, against approximately 33 percent from the aggregate score (main text Fig. 5D). Two controls confirm the effect reflects real structure rather than an artifact of contrast selection: permuting the mechanism labels collapsed accuracy to the 33 percent chance level, and listeners simulated with mixed mechanisms were assigned to one of their two constituent mechanisms rather than to the unrelated third.

### 8 Supplementary Figures

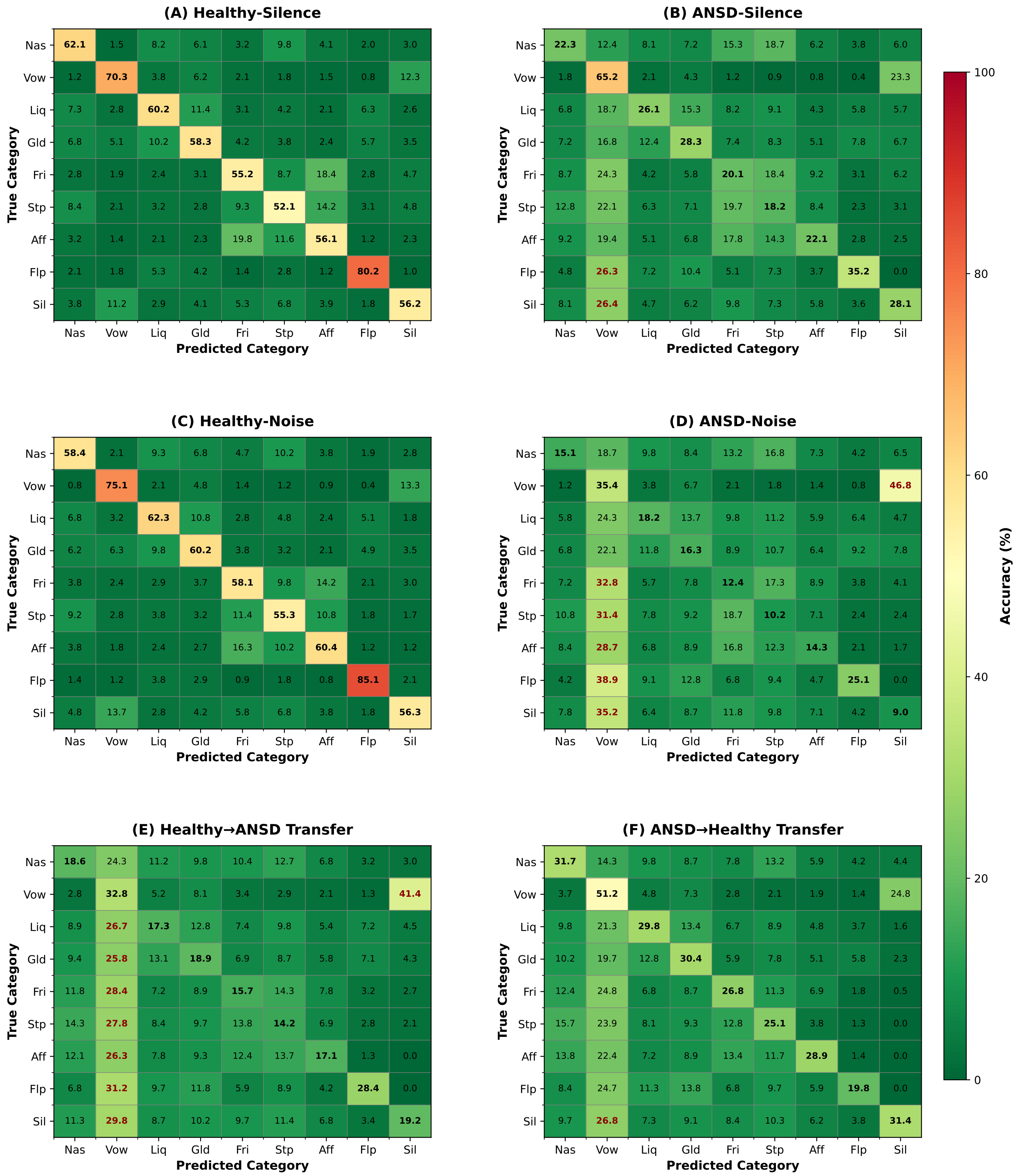

Figure S1: **Confusion matrices across conditions.** (A) Healthy-Silence, (B) ANSD-Silence, (C) Healthy-Noise, (D) ANSD-Noise, (E) Healthy→ANSD transfer, (F) ANSD→Healthy transfer. Key patterns: flaps drop from 80.2% (Healthy-Silence) to 35.2% (ANSD-Silence); vowel→silence confusion increases from 12.3% to 46.8% under ANSD-Noise.

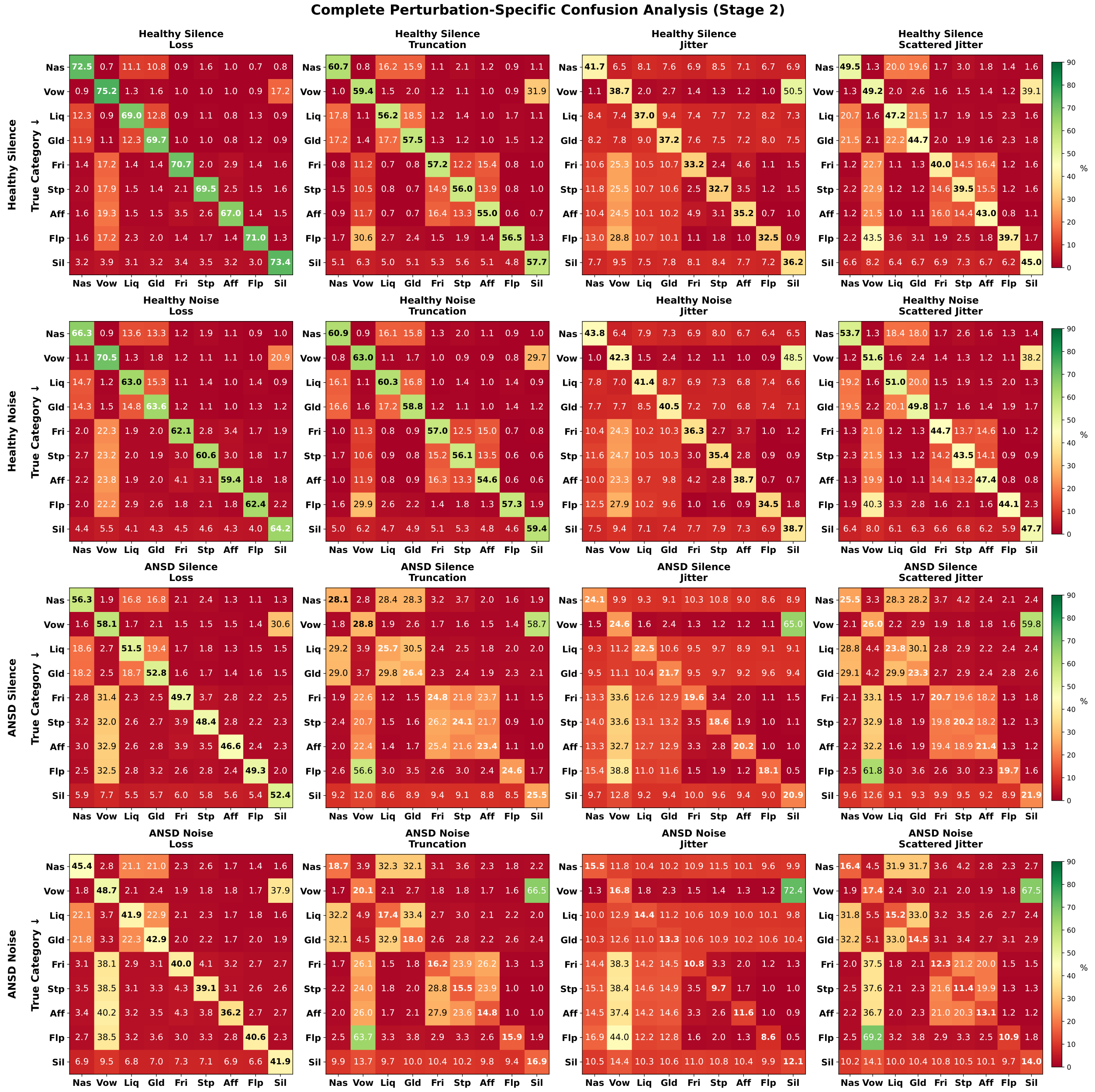

Figure S2: **Perturbation-specific confusion matrices.** Complete Stage 2 confusion matrices for all four trained models tested on individual perturbation types. Loss maintains temporal structure (within-manner confusions 11–18%). Truncation causes obstruent chain confusions and severe vowel→silence (32–67%). Jitter produces highest consonant→vowel (stop→vowel: 25–38%, flap→vowel: 31–44%). Scattered jitter shows intermediate patterns.

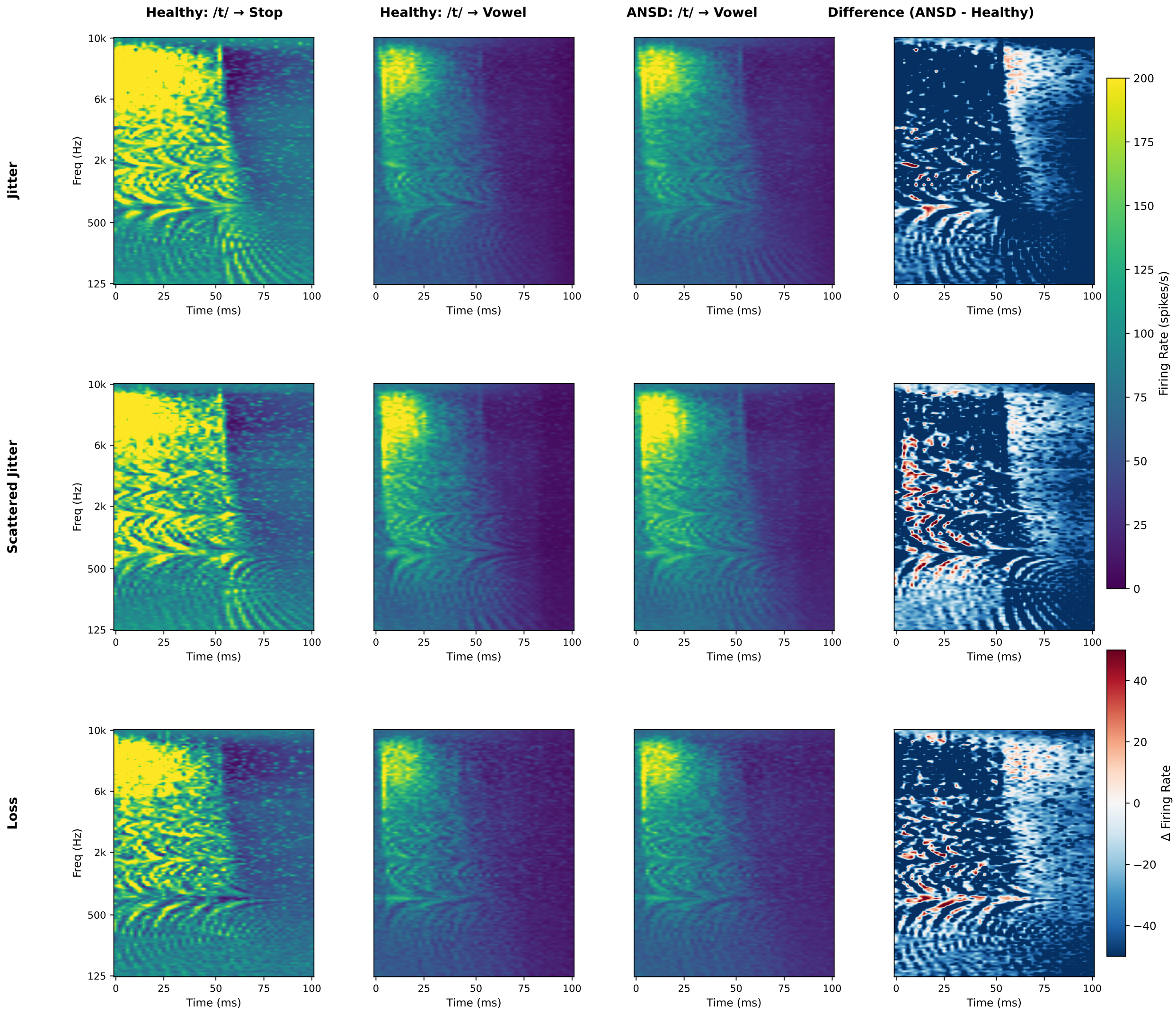

Figure S3: **Stop /t/ transformation across perturbation types.** Spectrotemporal templates for jitter (top), scattered jitter (middle), loss (bottom). Each row: (1) Healthy correct, (2) Healthy→vowel confusion, (3) ANSD→vowel confusion, (4) Difference map. Consistent pattern: reduced early burst (0–50 ms, blue), increased sustained activity (50–100 ms, red). Scattered jitter shows strongest effects. Truncation omitted (insufficient healthy correct templates, N<10).

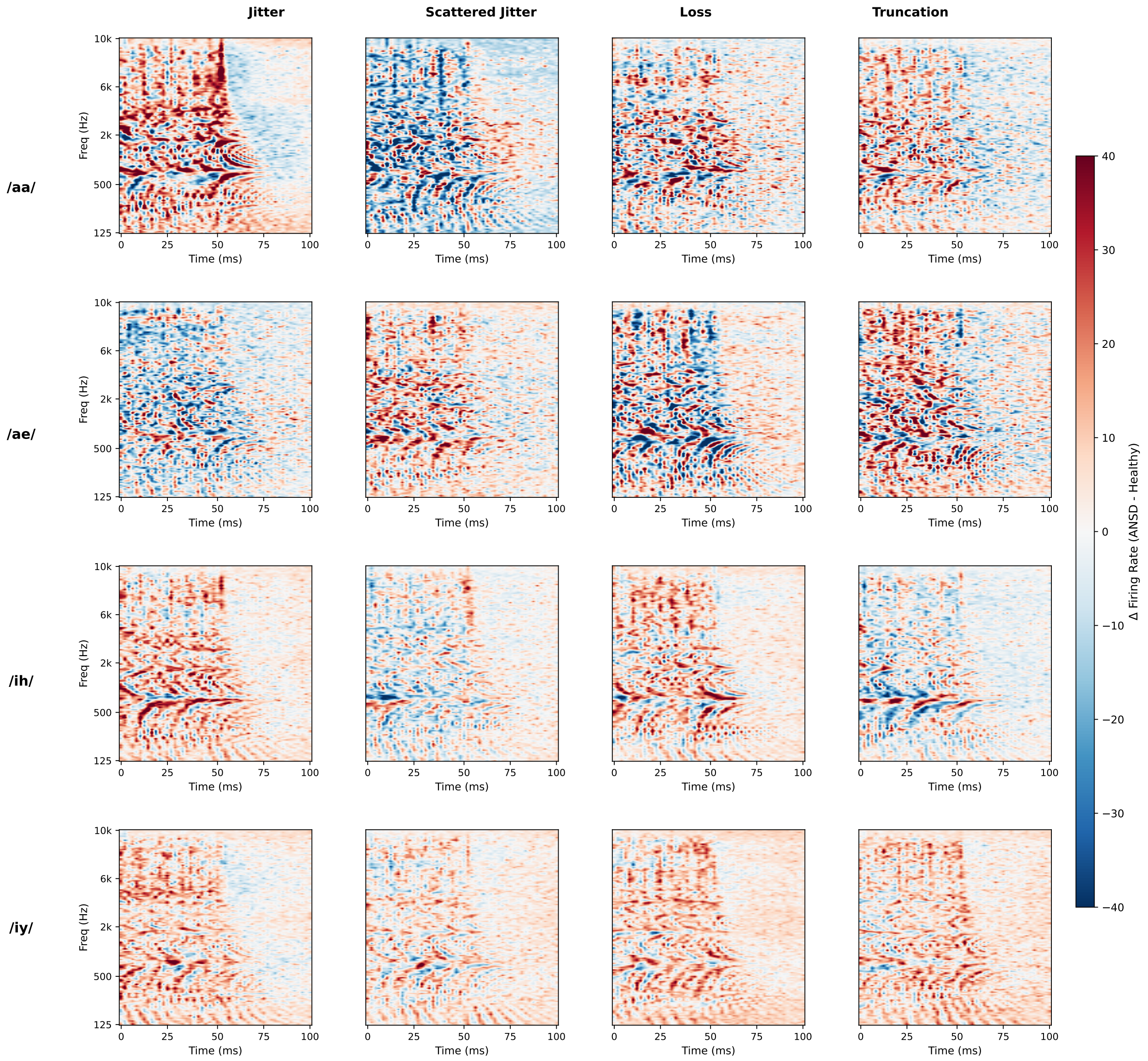

Figure S4: **Vowel encoding across perturbations.** Difference maps (ANSD−Healthy) for correctly classified vowels. Columns: jitter, scattered jitter, loss, truncation. Rows: /aa/, /ae/, /ih/, /iy/. Consistent patterns: high-frequency excess (6–10 kHz, 0–50 ms, red), mid-frequency reduction (500–2000 Hz, 25–75 ms, blue). Despite distortions, sufficient spectral structure preserved for classification (65.2% ANSD-Silence).

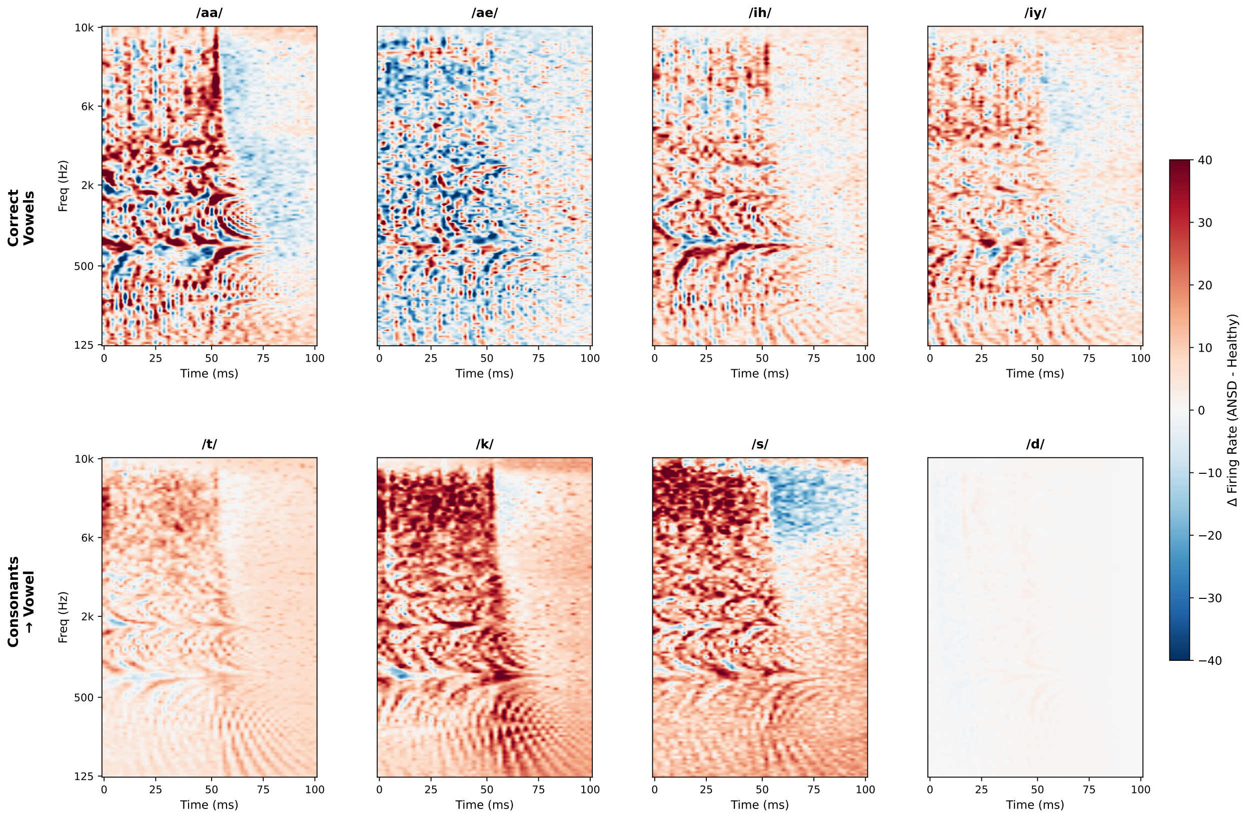

Figure S5: **Complete phoneme patterns (jitter).** Difference maps across phonetic inventory. Top: correctly classified vowels showing frequency/time-specific distortions. Bottom: consonants misclassified as vowels (/t/, /k/, /s/, /d/) showing transformation from brief transients to sustained patterns. Generality demonstrates fundamental spectrotemporal disruption rather than phoneme-specific effects.

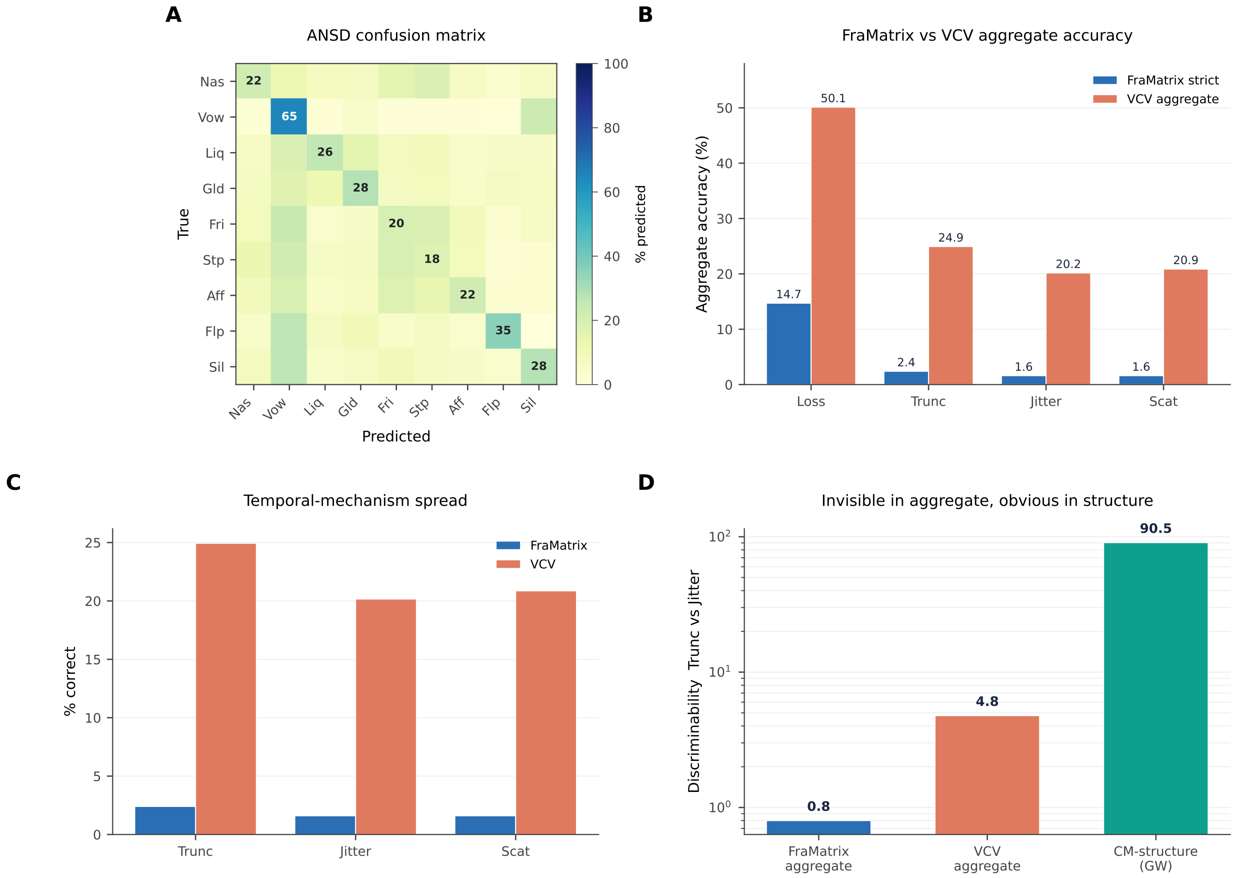

Figure S6: **The aggregate-scoring problem generalizes to an open-set format. (A)** Reference ANSD confusion-matrix structure (Stage 2). **(B)** Aggregate accuracy under the closed-set FraMatrix (strict) and the open-set VCV format, for each mechanism: open-set scores are higher across the board, but the temporal mechanisms still cluster. **(C)** Spread among the three temporal mechanisms (Truncation, Jitter, Scattered) stays small in both formats (0.8 pp FraMatrix, 4.8 pp VCV). **(D)** Truncation-versus-Jitter discriminability: negligible from either aggregate (0.8, 4.8) but large from confusion-matrix structure (90.5, Frobenius on the percentage-point scale). Structure separates the mechanisms far more than any aggregate, in both test formats.

[7] Gordon Wichern, Joe Antognini, Michael Flynn, Licheng Richard Zhu, Emmett McQuinn, Dwight Crow, Ethan Manilow, and Jonathan Le Roux. Wham!: Extending speech separation to noisy environments. *arXiv preprint arXiv:1907.01160*, 2019.
